# Targeting the Microbiome to Improve Gut Health and Breathing Function After Spinal Cord Injury

**DOI:** 10.1101/2023.06.23.546264

**Authors:** Jessica N. Wilson, Kristina A. Kigerl, Michael D. Sunshine, Chase E. Taylor, Sydney L. Speed, Breanna C. Rose, Chris M. Calulot, Brittany E. Dong, Tara R. Hawkinson, Harrison A. Clarke, Adam D. Bachstetter, Christopher M. Waters, Ramon C. Sun, Phillip G. Popovich, Warren J. Alilain

## Abstract

Spinal cord injury (SCI) is a devastating condition characterized by impaired motor and sensory function, as well as internal organ pathology and dysfunction. This internal organ dysfunction, particularly gastrointestinal (GI) complications, and neurogenic bowel, can reduce the quality of life of individuals with an SCI and potentially hinder their recovery. The gut microbiome impacts various central nervous system functions and has been linked to a number of health and disease states. An imbalance of the gut microbiome, i.e., gut dysbiosis, contributes to neurological disease and may influence recovery and repair processes after SCI. Here we examine the impact of high cervical SCI on the gut microbiome and find that transient gut dysbiosis with persistent gut pathology develops after SCI. Importantly, probiotic treatment improves gut health and respiratory motor function measured through whole-body plethysmography. Concurrent with these improvements was a systemic decrease in the cytokine tumor necrosis factor-alpha and an increase in neurite sprouting and regenerative potential of neurons. Collectively, these data reveal the gut microbiome as an important therapeutic target to improve visceral organ health and respiratory motor recovery after SCI.

**Research Highlights:**

- Cervical spinal cord injury (SCI) causes transient gut dysbiosis and persistent gastrointestinal (GI) pathology.
- Treatment with probiotics after SCI leads to a healthier GI tract and improved respiratory motor recovery.
- Probiotic treatment decreases systemic tumor necrosis factor-alpha and increases the potential for sprouting and regeneration of neurons after SCI.
- The gut microbiome is a valid target to improve motor function and secondary visceral health after SCI.

## INTRODUCTION

The gut-central nervous system (CNS) axis is a bi-directional pathway where disruptions in the normal fauna and flora of the gut, i.e. gut dysbiosis, can impair CNS function. Conversely, CNS injury or disease can lead to gut dysbiosis. As a result of this unique interrelationship, gut dysbiosis has been linked to a variety of neurological disorders, including depression, amyotrophic lateral sclerosis, and cognitive decline or impairment ^1–3^. In this study, we investigated how cervical spinal cord injury can lead to gut dysbiosis, which can potentially further lead to greater CNS dysfunction and impaired functional recovery of breathing (Figure 1 a; Schematic).

**Figure 1.**
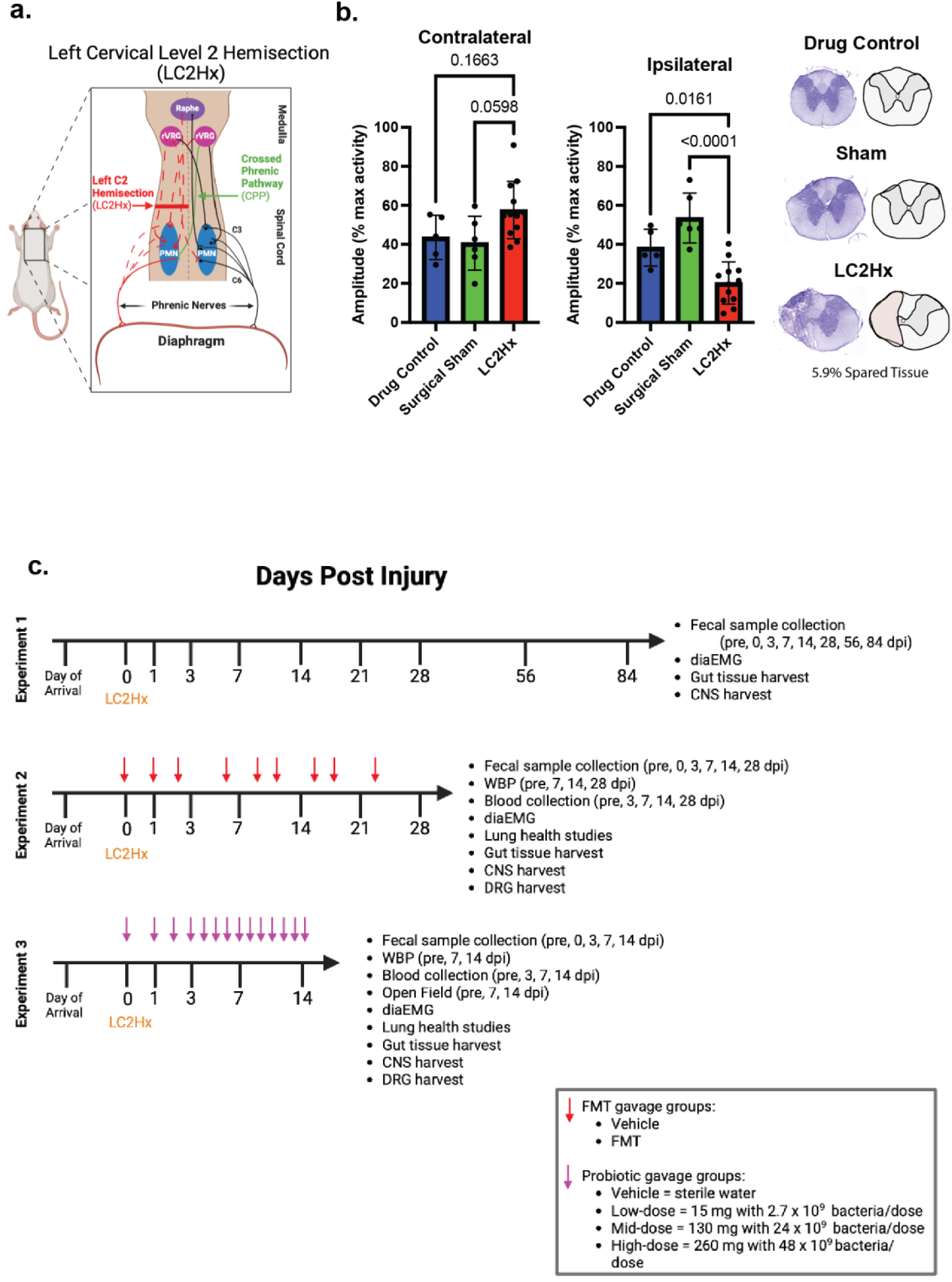
Injury model and experimental design. **a**, Schematic representation of the LC2Hx model disrupting the respiratory pathway. **b**, Quantification of diaphragm burst amplitude as a percent maximum activity contralateral and ipsilateral to the LC2Hx and histological images indicating the injury severity for each group (blue = drug control (n = 5), green = sham (n = 6), red = LC2Hx (n = 12)). As expected, the LC2hx produced a significant decrease in the burst amplitude on the ipsilateral side of the lesion (One-way ANOVA p < 0.0001 with Tukey’s posthoc LC2Hx/drug control p = 0.0161, LC2Hx/ surgical sham p < 0.0001). **c.** Experimental design; Experiment 1: Evaluating gut dysbiosis following LC2Hx, groups: drug control (n = 5), surgical sham (n = 6), LC2hx (n = 12); Experiment 2: Fecal matter transplant (FMT) treatment following LC2Hx, groups: vehicle (FMT slurry without fecal component, n = 12) or FMT-treated (n = 12), three consecutive gavages beginning day of surgery followed by biweekly gavage until endpoint; Experiment 3: Probiotic treatment following LC2Hx, groups: vehicle (sterile water, n = 5), low-dose probiotic = 15 mg with 2.7 x 10^9^ bacteria/dose (n = 6), mid-dose probiotic = 24 x 10^9^ bacteria/dose (n = 6), high-dose probiotic = 24 x 10^9^ bacteria/dose (n = 5) gavage daily beginning day of injury.

## RESULTS

### Cervical SCI leads to acute gut dysbiosis and gastrointestinal tract pathology

Nearly 60% of all SCI cases are at the cervical level. To model this type of injury, we used a well-characterized left C2 hemisection (LC2Hx) model of SCI in rats. This lesion reduces diaphragm activity and impairs ventilation (Figure 1 a-b) ^4, 5^. In Experiment 1 (Figure 1c), adult female Sprague Dawley rats were randomized into three experimental groups: 1) LC2Hx (n = 12), 2) surgical sham (n = 6; received post-surgical drugs and all surgical procedures except spinal hemisection), and 3) drug control (n = 5; no surgical procedure but animals were anesthetized and given post-surgical drugs) (Figure 1 c; Experimental design). Because anesthesia and opioid-based analgesics affect normal gut function and the gut microbiome, a surgical drug group was a necessary control ^6, 7^. To prevent bacterial transfer via coprophagia, animals were co-housed throughout the experiment and cage-mates were in the same experimental group.

Electromyography (EMG) of the diaphragm ipsilateral to the injury was significantly lower in LC2Hx animals compared to surgical sham and drug control animals (Figure 1 b; One-way ANOVA p < 0.0001, F = 18.77 with Tukey’s posthoc LC2Hx/drug control p = 0.0161; LC2Hx/surgical sham p < 0.0001). To determine if LC2Hx affects the gut microbiome, fecal samples were collected from each animal of same-group co-housed rats pre-injury (i.e., 0 days post-injury (dpi)) and on days 1, 3, 7, 14, 21, 28, 58, and 84 dpi then were prepared for 16S rRNA sequencing of hypervariable region V4 (using 515F-806R primers). Following 16S rRNA sequencing, we performed principal component analysis on the Bray Curtis distances of normalized operational taxonomic units rarefied reads. There was a significant separation of the samples from the surgical sham and LC2Hx groups in the ordination space at the operational taxonomic unit level (Figure 2 a; PERMANOVA p = 0.001). Indeed, 16S rRNA analysis of fecal samples prior to and following LC2Hx SCI revealed a rapid and robust change in the composition of the gut microbiome. Notably, the Firmicutes to Bacteroidetes (F:B) ratio increased significantly at one-day post-injury in LC2Hx but not drug control or surgical sham animals (Figure 2 b; Two-way ANOVA time x treatment p < 0.0011; F_(18)_ = 2.44). At the phylum level, Firmicutes and Bacteroidetes bacteria constitute 90% of the gut microbiome ^8, 9^. An increase in the F:B ratio has been used previously to describe gut dysbiosis, or an imbalance of pathogenic and beneficial bacteria ^10^. Further, a high F:B ratio has been associated with increased susceptibility to inflammation and obesity ^10–13^. The presence of a high F:B ratio following SCI has also been shown previously in thoracic-level mouse injury models ^14^. However, this finding has been inconsistent among other preclinical models of SCI ^14–16^.

**Figure 2.**
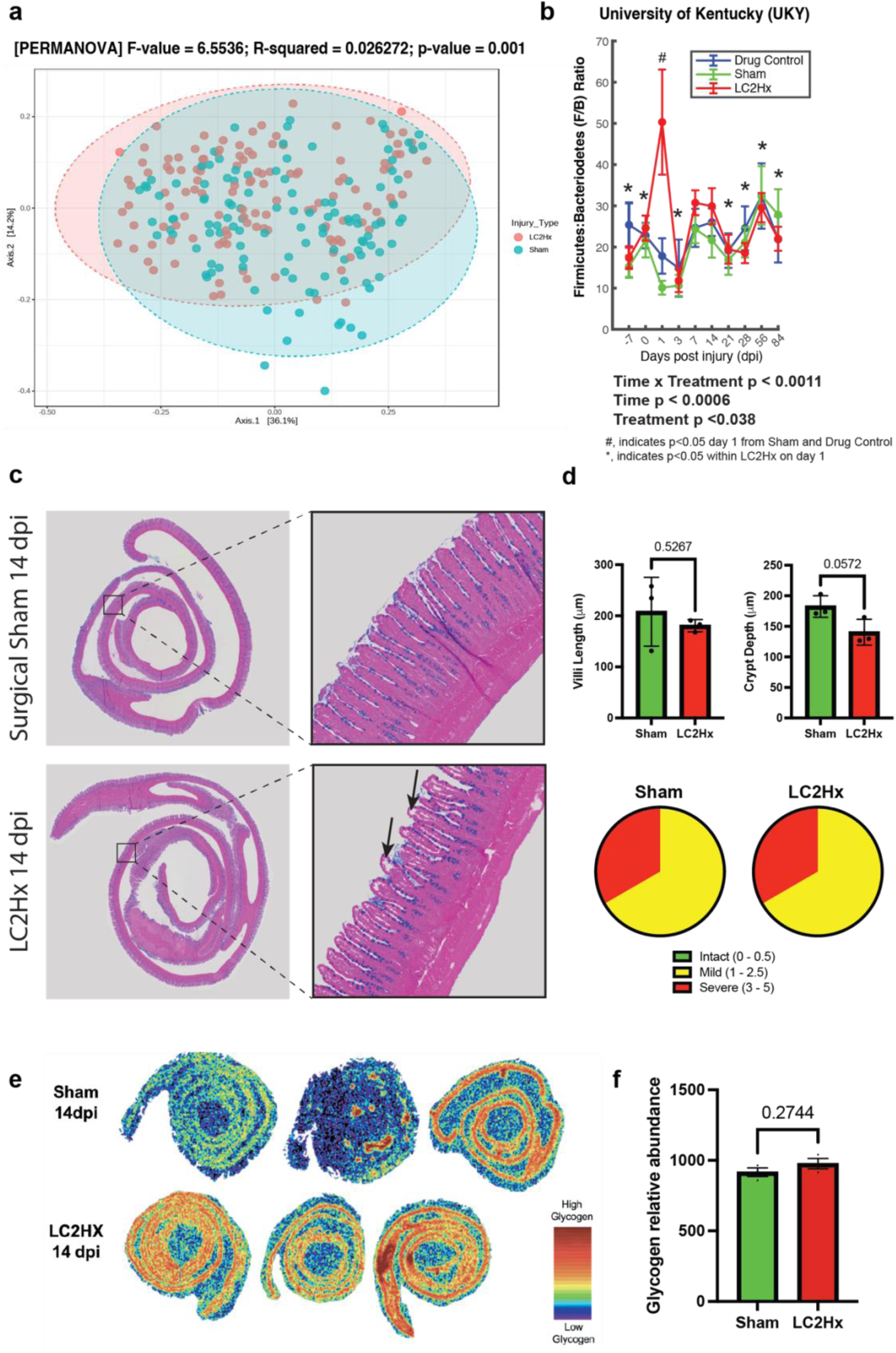
Cervical SCI leads to acute dysbiosis and GI tract pathology. **a,** Principal coordinate analysis (PCoA) on Bray Curtis distances of normalized operational taxonomic units (OTUs) rarefied reads colored by day post-injury in surgical sham (square) or LC2Hx (circle) (permnova p = 0.0001). **b,** 16S rRNA sequencing data revealed a significant increase in the Firmicutes:Bacteriodetes ratio at 1 dpi in the LC2hx group (Two-way RM ANOVA p < 0.0011). **c - d**, Histological representations of ileal tissue pathology at 14 dpi revealed no change in villi length, although there was a strong trend in the reduction of crypt depth (n = 3/group; unpaired t-test, two-tailed; villi length p = 0.52; crypt depth p = 0.058). Pie charts representing pathological changes in ileal tissue utilizing the Chiu/Park scoring system in the sham versus LC2Hx animals at 14 days post-LC2Hx revealed no differences between groups, although epithelial lifting, an indication of damage marked by black arrows, was evident in the LC2Hx group (0 – 0.5 = intact (green); 1.0 – 2.5 = mild (yellow); 3 – 5 = severe (red)). **e**, MALDI imaging revealed that LC2hx leads to metabolic dysregulation of glycogen in the ileum at 14 dpi; *M/z* 1013.3213 (6 linear chains of glucose) displayed as representative image of glycogen. **f,** Quantification of MALDI imaging (n = 3/group; unpaired, two-tailed t-test; p = 0.2744; R^2^ = 0.285). Bars represent mean ± SD. Bolded text indicates a p-value < 0.05.

Although the F:B ratio returned toward pre-injury levels by three days post-injury, histological evaluation of the gastrointestinal (GI) tract within the ileum revealed persistent and profound gut pathology. Specifically, subepithelial blebbing and disruption of the epithelial lining were consistent in the gut of LC2Hx SCI rats at 14 dpi (Figure 2 c. – marked by black arrows). Subepithelial blebs found at the tips of villi, called Gruenhagen spaces ^17, 18^, are an indication of ischemia within the intestine and can lead to disruption and disintegration of the epithelial lining if ischemia is prolonged ^19^. To quantify this observed damage, the Chiu/Park scoring system of GI tract pathology, which grades the degree of intestinal mucosal injury on a scale of 0 – 5 (0 = normal mucosa; 1 = subepithelial space at the villous tip; 2 = extension of subepithelial space with moderate lifting; 3 – massive lifting down the sides of villi, some denuded tips; 4 = denuded villi, dilated capillaries; 5 = disintegration of lamina propria) ^20^. Blinded evaluation of the tissue using the Chiu/Park scoring system of GI tract pathology revealed no differences in the pathology and intestinal injury in sham versus LC2Hx animals, although epithelial lifting, an indication of damage marked by black arrows, was evident in LC2Hx animals (Figure 2 c - d; representative pie charts utilizing the Chiu/Park score). Further, neurogenic shock and autonomic nervous system impairment following SCI can cause hypoperfusion of visceral organs, inducing inflammation and pathological changes ^21, 22^. This hypoperfusion and loss of epithelium can induce further mucosal atrophy through necrosis and apoptosis of crypt cells. Since crypt cells provide stem cells that develop and populate the villi, crypt destruction can result in decreased villi length ^23^. We observed no change in villi length in animals 14 days post L2CHx (Figure 2 d; unpaired, two-tailed t-test; p = 0.5267; t = 0.6925); however, there was a strong trend in the reduction of crypt depth, although this change was not statistically significant (Figure 2 d; unpaired, two-tailed t-test; p = 0.0572; t = 2.646).

Utilizing matrix-assisted laser desorption ionization (MALDI) imaging, we found slightly increased levels of glycogen in the ileal tissue of LC2Hx animals compared to the surgical sham, although this increase was not statistically significant (Figure 2 e-f; unpaired, two-tailed t-test, p = 0.2744; t = 1.265; R^2^ = 0.2859). Glycogen is a storage macromolecule that can influence bacterial survival during times of extreme stress ^24^. Glycogen stores allow for protein N-glycosylation in the brain, and dysregulation can cause disease ^25, 26^. Additionally, glycosylation is a hallmark of intestinal mucins, or glycoproteins, that act as the first line of defense to the intestinal epithelium against pathogens and luminal microbiota ^27^. Interestingly, microbiota have been found to regulate host protein glycosylation, which has been linked to intestinal health and disease ^28^. This relationship further emphasizes the importance of maintaining a healthy gut microbiome. The increasing glycogen levels in ileal tissue 14 days after LC2Hx SCI, may be a compensatory response to increase mucin production and repair the intestinal barrier in response to the increase in the F:B ratio that occurs immediately after SCI.

These findings expand upon previous observations of the ileum at 24 hours post-injury at the thoracic level (T7/8 transection), where the proliferation of submucosal lymphatic tissue and widening of mucosal epithelial cell interspaces were apparent ^29^. Our findings are also consistent with previous findings in low thoracic injuries (T10/11 contusion), where decreased mucosal layer thickness and persistent atrophy began at 48 hours post-injury and were evident until 4 weeks post-injury ^30^. A separate T10 contusion study described similar findings of disruption in the jejunum at 3dpi, with evidence of decreased villi height, increased crypt depth, and increased subepithelial space at the tip of villi ^31^. Our results are the first to show that the high cervical LC2Hx model can provide meaningful insights into the serious problem of bowel and GI tract disease and pathology after SCI. Indeed, gastrointestinal complications following SCI are a major factor impacting quality of life and account for 11% of rehospitalizations in the first year post-SCI ^32, 33^. Furthermore, individuals with injuries occurring at the cervical level are more likely to experience constipation and abdominal bloating and are more likely to require re-hospitalization relative to individuals with lower-level injuries ^34, 35^. Gastrointestinal dysfunction also can compromise intestinal barrier function, promote bacterial translocation in which intestinal bacteria leak into the bloodstream, and contribute to systemic inflammation and multiple organ dysfunction syndrome (MODS), further impacting recovery and quality of life of SCI individuals ^36^.

### Targeting gut dysbiosis improves ventilatory function and gut health

Since gut dysbiosis and persistent GI tract pathology are evident after SCI, we aimed to test whether oral gavage of either fecal matter transplant (from young, uninjured animals) or clinical-grade probiotics would restore the gut and microbiome to their pre-injury state. Indeed, previous studies have demonstrated that gut dysbiosis can induce systemic inflammation ^37–39^, and separately, it has been shown that systemic inflammation can impair respiratory plasticity40,41.

Additionally, it has been established that treating neurotrauma-induced gut dysbiosis leads to improved locomotor recovery, smaller lesion sizes, and prevention of anxiety-like behavior ^14, 42^. However, the impact of treating gut dysbiosis following LC2Hx on breathing function, GI tract pathology, and its long-term effects remain unexplored. We aimed to answer these questions and hypothesized that the dynamic changes in the gut microbiome immediately post-LC2Hx impede functional recovery of breathing in the acute to the sub-acute phase of SCI. Further, we hypothesized that treatment of the resulting neurotrauma-induced gut dysbiosis would improve gut health and allow for a more profound recovery of breathing at acute time points post-injury.

Animals received fecal matter transplants (FMT) for three consecutive days, beginning the day of injury, followed by bi-weekly gavage until the predetermined endpoint. This gavage regimen combines two prior reports of FMT studies in rats (Schmidt et al., 2020, Kelly et al., 2016) ^3, 42^. Treating LC2Hx animals with FMT derived from young, uninjured animals led to marginal improvements in tidal volume in the first cohort of animals (Supplemental Figure 1 a; n = 4/group Two-way ANOVA time x treatment p = 0.2666; F_(3, 18)_ = 1.431). To differentiate whether this change was due to alterations in peripheral lung mechanics or recovery in central respiratory pathways, we assessed lung mechanics using the flexiVent system. There was an increased inspiratory capacity in FMT-treated animals, which may indicate that the FMT treatment mitigated atelectasis, or partial lung collapse (Supplemental Figure 1 b; Welch’s unpaired, two-tailed t-test; inspiratory capacity p = 0.0585; t = 2.579). Additional measures of lung function, such as dynamic compliance, dynamic resistance, and bronchioalveolar lavage (BAL) cells, did not differ between groups suggesting the overall lung mechanic findings did not account for the increase in tidal volume. Therefore, these findings suggest that the change in tidal volume may be driven centrally rather than by peripheral lung mechanic compensation. However, when all cohorts were completed (n = 12/group), this observation of improved tidal volume disappeared, and there were no statistically significant differences between groups in whole-body plethysmography (WBP) or diaphragmatic EMG (diaEMG) measurements (Figure 3 a and b). Inspiratory capacity remained elevated with FMT treatment, although not significantly, while dynamic compliance, dynamic resistance, and BAL cells remain unchanged between groups (Supplemental Figure 1 c; Welch’s unpaired, two-tailed t-test; inspiratory capacity p = 0.1122; t = 1.719).

**Figure 3.**
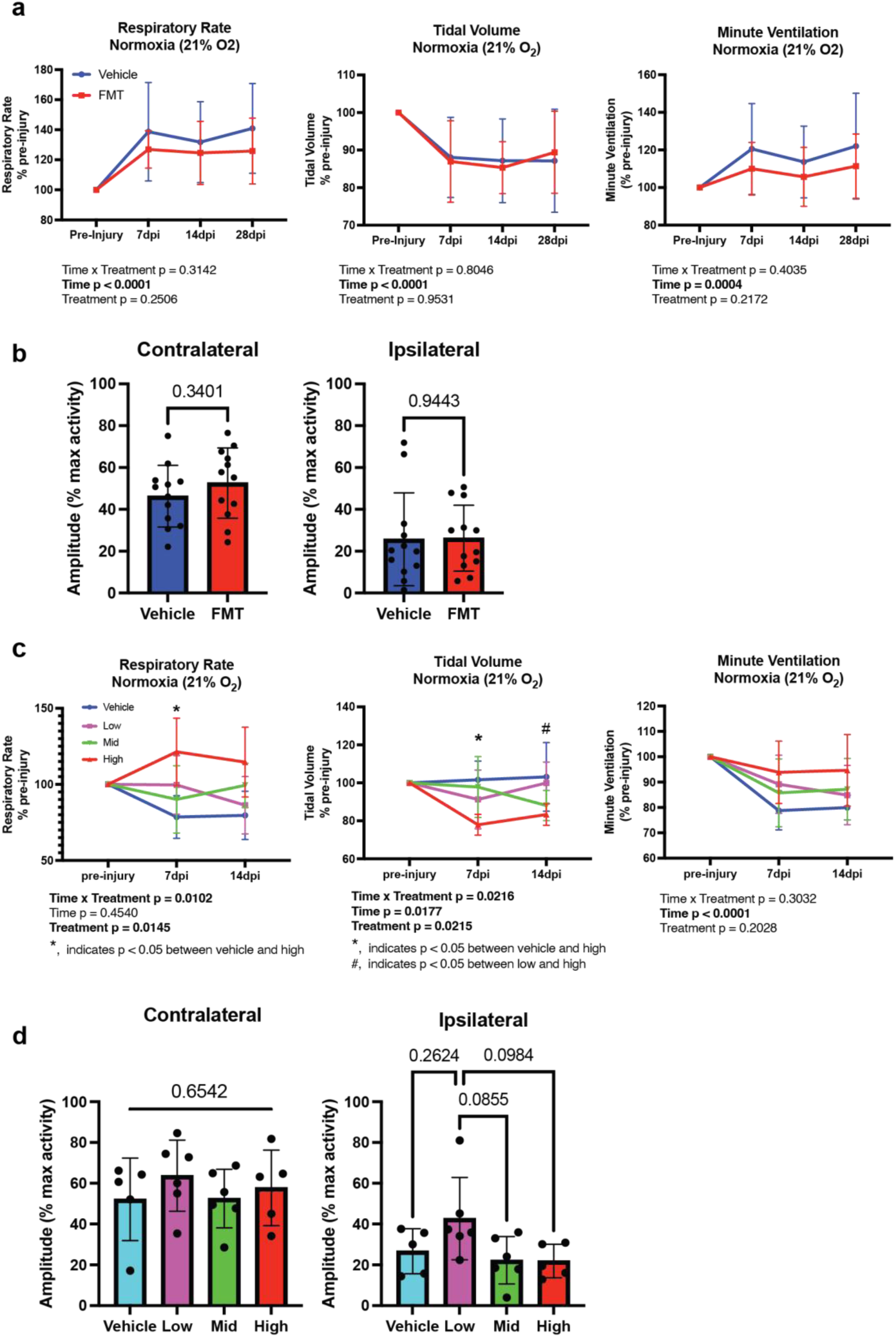
Targeting gut dysbiosis improves ventilatory function. **A,** Quantification of WBP measurements (respiratory rate, tidal volume, and minute ventilation) and **b,** diaEMG studies following LC2Hx and treatment with vehicle (blue; n = 12) or FMT (red; n = 12) revealed marginal differences between groups. c – d, Quantification of WBP and diaEMG studies following LC2Hx and treatment with vehicle (blue; n = 5), low-dose (purple; n = 6), mid-dose (green; n = 6), or high-dose (red; n = 5) probiotics. **c,** Treatment with probiotics following LC2Hx revealed a significant increase in the respiratory rate in the high-dose probiotics versus vehicle-treated animals at 7 dpi (Two-way RM ANOVA with Tukey’s posthoc vehicle/high 7dpi p = 0.0340). Conversely, there was a significant decrease in tidal volume in the high-dose probiotic versus vehicle-treated animals at 7 dpi and high-dose versus low-dose probiotic-treated animals at 14 dpi (Two-way RM ANOVA with Tukey’s posthoc high/vehicle 7 dpi p = 0.0121 and high/low 14 dpi p = 0.0496) Although not significantly different, all probiotic treated animals were able to maintain a higher minute ventilation than vehicle-treated animals at all time points post-injury. **d,** diaEMG studies revealed no significant differences between any treatment groups in the contralateral or ipsilateral diaphragm following LC2Hx, although the low-dose probiotic treated group was able to mount a greater burst amplitude compared to all other groups (One-way ANOVA with Tukey’s posthoc low/vehicle p = 0.2624, low/mid p = 0.0855, low/high p = 0.0984). Bars represent mean ± SD. Bolded text indicates a p-value < 0.05.

As FMT treatment led to mixed and inconsistent improvements, we initiated an alternative strategy utilizing a standardized probiotic treatment to mitigate gut dysbiosis based on previous findings (Kigerl et al., 2016). We implemented a daily probiotic regimen consisting of a low, mid, and high dose, as the literature describing dosage of probiotics given to rats was highly variable ^43–46^. The gavage volume was maintained at 0.5 ml, but the bacterial concentration increased from 2.7 x 10^9^, to 24 x 10^9^, to 48 x 10^9^ bacteria/dose in our rat LC2Hx model. In WBP studies under normoxic conditions (21% O_2_), animals treated with high-dose probiotics showed the expected reduction in tidal volume but were able to increase their respiratory rate at 7 dpi compared to the vehicle-treated group (Figure 3 c. Two-way RM ANOVA tidal volume time x treatment p = 0.0216; F (6, 36) = 2.875; frequency time x treatment p = 0.0102; F (6, 36) = 3.341with Tukey’s posthoc; tidal volume vehicle/high p = 0.0074; frequency vehicle/high p = 0.0210). Minute ventilation was maintained in the high-dose group at 7 dpi and 14 dpi due to this compensatory increase in the respiratory rate (Figure 3 c; Two-way RM ANOVA minute ventilation time x treatment p = 0.3032; F (6. 36) = 1.253 with Tukey’s posthoc; vehicle/high 7dpi p = 0.1221; vehicle/high 14 dpi p = 0.1712). Interestingly, the vehicle, low-dose and mid-dose animals did not display the expected SCI-dependent increase in respiratory rate and decrease in tidal volume. To investigate this, animals were challenged under hypoxic conditions (11% O_2_). we began to see the expected SCI-dependent increase in respiratory rate, decrease in tidal volume and compensation of minute ventilation in all probiotic treated animals (low-, mid-, and high-dose) at 7 dpi. Interestingly, minute ventilation was increased at 7dpi in probiotic treated animals when compared to vehicle-treated animals with the mid-dose group mounting the highest response (Supplemental Figure 2 a; Two-way RM ANOVA with Tukey’s posthoc vehicle/mid 7dpi p = 0.1864). Examinations of lung function indicated no statistically significant differences in lung compliance, resistance, or BAL cell counts (Supplemental Figure 2 b.). However, the inspiratory capacity for the high-dose group was greater than all other groups and reached significance when compared to the low-dose group (Supplemental Figure 2 b; One-way ANOVA p = 0.0385; F = 3.452 with Tukey’s posthoc high/low p = 0.0239). Interestingly, when we evaluated diaphragm EMG, the animals in the low-dose group were able to produce the largest burst amplitude ipsilateral to the injury compared to all other groups (Figure 3 d; One-way ANOVA p = 0.0635; F = 2.898 with Tukey’s posthoc low/vehicle p = 0.2624; low/mid p = 0.0855; low/high = 0.0984). Importantly, there were no differences in injury severity between groups based on spared tissue analysis. This indicates that these improvements in respiratory function were not due to a less severe injury in any particular group (Supplemental Figure 3 a – b. One-way ANOVA p = 0.9411; F = 0.1298). While probiotic treatment improved respiratory motor function, we did not observe changes in overall locomotor function demonstrated by the open field test (Supplemental Figure 4). These findings suggest that probiotic treatment following cervical spinal cord injury may improve respiratory recovery through compensatory mechanisms by increasing the respiratory rate and amplitude of the diaphragmatic output.

In addition to improvements in ventilatory function, treatment with probiotics following SCI also led to improved gut pathology within the ileal tissue of the small intestine. FMT treatment produced marginal improvement in gut pathology. Less gaping between villi was evident with FMT treatment, an indication of a healthier physiological state within the gut as these projections are important for nutrient absorption via increased surface area of the small intestine. There were minimal increases in villi length, although no differences in crypt depth or Chiu/Park scores, indicating a marginal improvement in gut pathology following FMT treatment compared to vehicle-treated animals (Figure 4 a – d; b. unpaired, two-tailed t-test villi length p = 0.2807; t = 1.107; c. unpaired, two-tailed t-test crypt depth p = 0.4801; t = 0.7189; d. representative pie charts utilizing the Chiu/Park score). Of note, the endpoint for the FMT study was 28 dpi in comparison to animals in Experiment 1, as well as animals in Experiment 3 (i.e., probiotic treatment), which both had an endpoint of 14 dpi. Because we observed the initial improvement in total volume by 14 dpi (Supplemental Figure 1 a) but no changes in gut pathology by 28 dpi in Experiment 2 (Figure 4 a – d), we used the 14 dpi timepoint for Experiment 3. The extra two weeks in Experiment 2 may have allowed more healing and repair making it difficult to detect differences with FMT treatment. Probiotic treatment resulted in increases in villi length in a dose-dependent manner, although these differences were not statistically significant, while there were no differences in crypt depth (Figure 4 e – h. One-way ANOVA; villi length p = 0.1817; F = 1.808; crypt depth p = 0.8036; F = 0.3302). Additionally, we observed an improvement in “intact” ileal tissue indicated by the Chiu/Park score with the mid and high-dose probiotic-treated animals compared to the low-dose probiotic and vehicle-treated animals. (Figure 4 h; representative pie charts utilizing the Chiu/Park score). Overall, these findings suggest that treatment of gut dysbiosis acutely after SCI improves gut pathology and potentially improves gut function.

**Figure 4.**
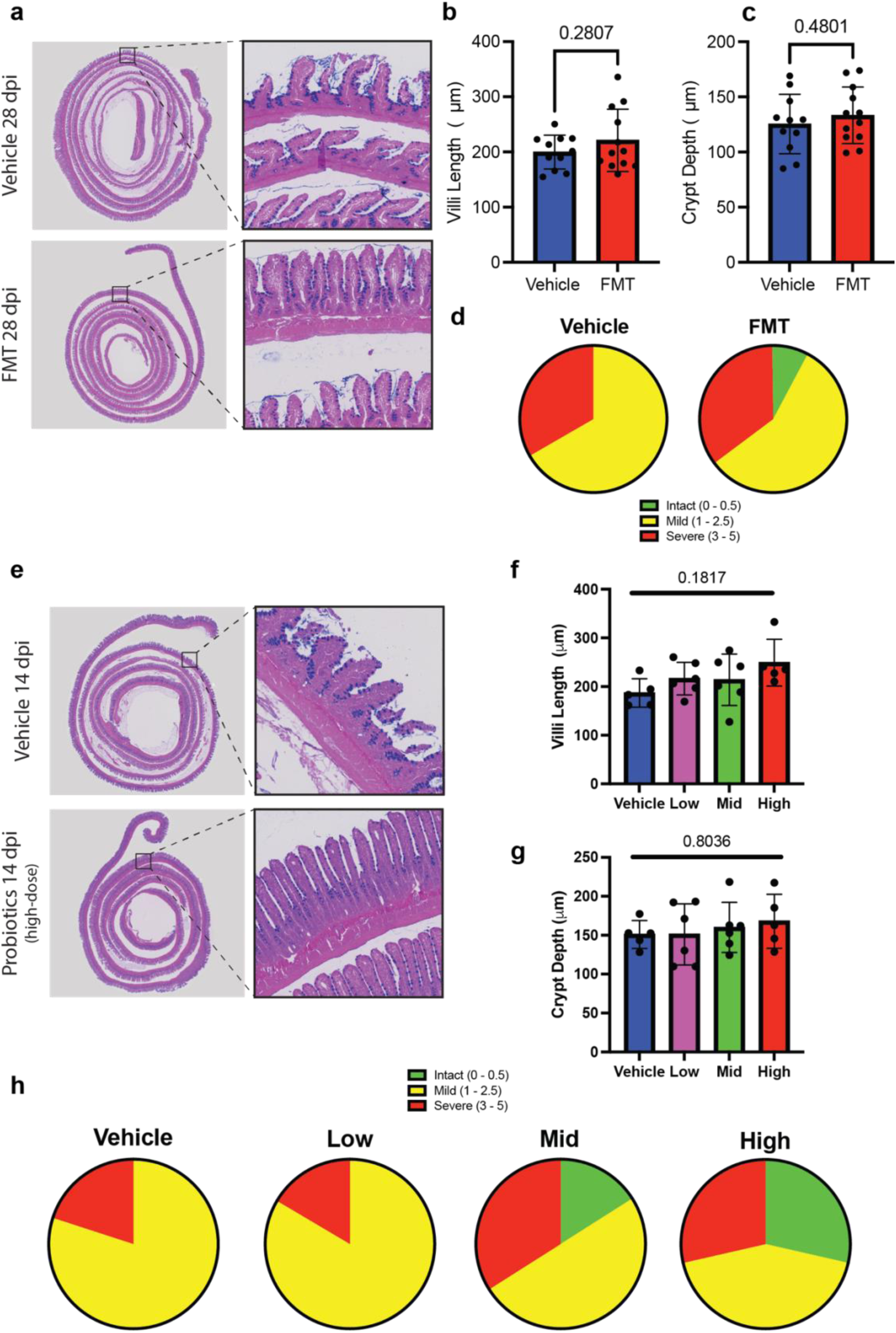
Targeting gut dysbiosis improves gut health. **a,** Representative histological images of ileal tissue stained with Alcian blue in vehicle (blue) and FMT-treated (red) animals at 28 days post-LC2Hx at 2x (left) and 5x (right - enlarged window). **b – d,** Quantification of **b,** villi length (unpaired t-test p = 0.2807), **c,** crypt depth (unpaired t-test p = 0.4801), and injury severity represented by the **d,** Pie charts representing pathological changes in ileal tissue utilizing the Chiu/Park score in the vehicle versus FMT-treated animals at 28 days post-LC2Hx revealed minimal changes between groups (0 – 0.5 = intact (green); 1.0 – 2.5 = mild (yellow); 3 – 5 = severe (red)). **e,** Representative histological images of ileal tissue stained with Alcian blue in vehicle and high-dose probiotic-treated animals at 14 days post-LC2Hx at 2x (left) and 5x (right - enlarged window). **f – h,** Quantification of ileal histology at 14 days post-LC2Hx in vehicle (blue), low-dose (purple), mid-dose (green), or high-dose (red) probiotic-treated animals. **f,** Quantification of villi length revealed a trend of increased length as the probiotic dose increased compared to vehicle, although these differences were not significant (One-way ANOVA with Dunnett’s posthoc p = 0.1817; vehicle/low p = 0.5316; vehicle/mid p = 0.5906; vehicle/high p = 0.0791). **g,** Quantification of crypt depth revealed no significant difference between groups (One-way ANOVA with Dunnett’s posthoc p = 0.8036; vehicle/low p > 0.999; vehicle/mid p = 0.9362; vehicle/high p = 0.7399). **h,** Pie charts representing pathological changes in ileal tissue utilizing the Chiu/Park score in the vehicle, low-, mid-, or high-dose probiotic-treated animals at 14 days post-LC2Hx revealed an improvement in pathology (0 – 0.5 = intact (green); 1.0 – 2.5 = mild (yellow); 3 – 5 = severe (red)). Bars represent mean ± SD. Bolded text indicates a p-value < 0.05.

### Probiotic treatment after SCI reduces systemic inflammatory responses and increases the sprouting and regenerative potential of neurons

To investigate the mechanism of these functional and pathological improvements, we obtained serum samples throughout the experiment to assess systemic inflammation. We observed a decrease in TNF-alpha at all time points in both FMT and probiotic-treated animals compared to vehicle-treated. This reduction was significant at 28 dpi in FMT-treated animals (Figure 5 a; Two-way RM ANOVA time x treatment p = 0.0448; F (4, 88) = 2.548 with Bonferroni posthoc 28 dpi p = 0.0062) and at 3 dpi in all probiotic-treated animals, regardless of dose level (Figure 5 b; Two-way RM ANOVA time x treatment p = 0.0489; F (9, 54) = 2.068 with Tukey’s posthoc 3dpi vehicle/low p = 0.0316, vehicle/mid p = 0.0152, vehicle/high p = 0.013). Additionally, there was an overall effect of FMT treatment on IL-6 serum levels, with FMT-treated animals having lower IL-6 compared to vehicle-treated (Figure 5 a; Two-way RM ANOVA treatment p = 0.0459; F (1, 22) = 4.478). These findings are consistent with clinical observations. The pro-inflammatory cytokine IL-6 was found to be elevated in patients at 48 hours post-injury but decreased by 7 days post-injury ^47^. Conversely, an alternate study revealed that this increase is persistent as levels of IL-6, as well as TNF-alpha, are significantly elevated levels in SCI patients in the post-acute (2-52 weeks post-injury) and chronic (>52 weeks post-injury) phases of recovery compared to healthy controls ^48^. Importantly for the recovery of breathing, systemic inflammation has been shown to impair respiratory plasticity ^41^. Therefore, the reduction in systemic inflammation following treatment with probiotics may play an important role in the improved minute ventilation observed in Figure 3 c.

**Figure 5.**
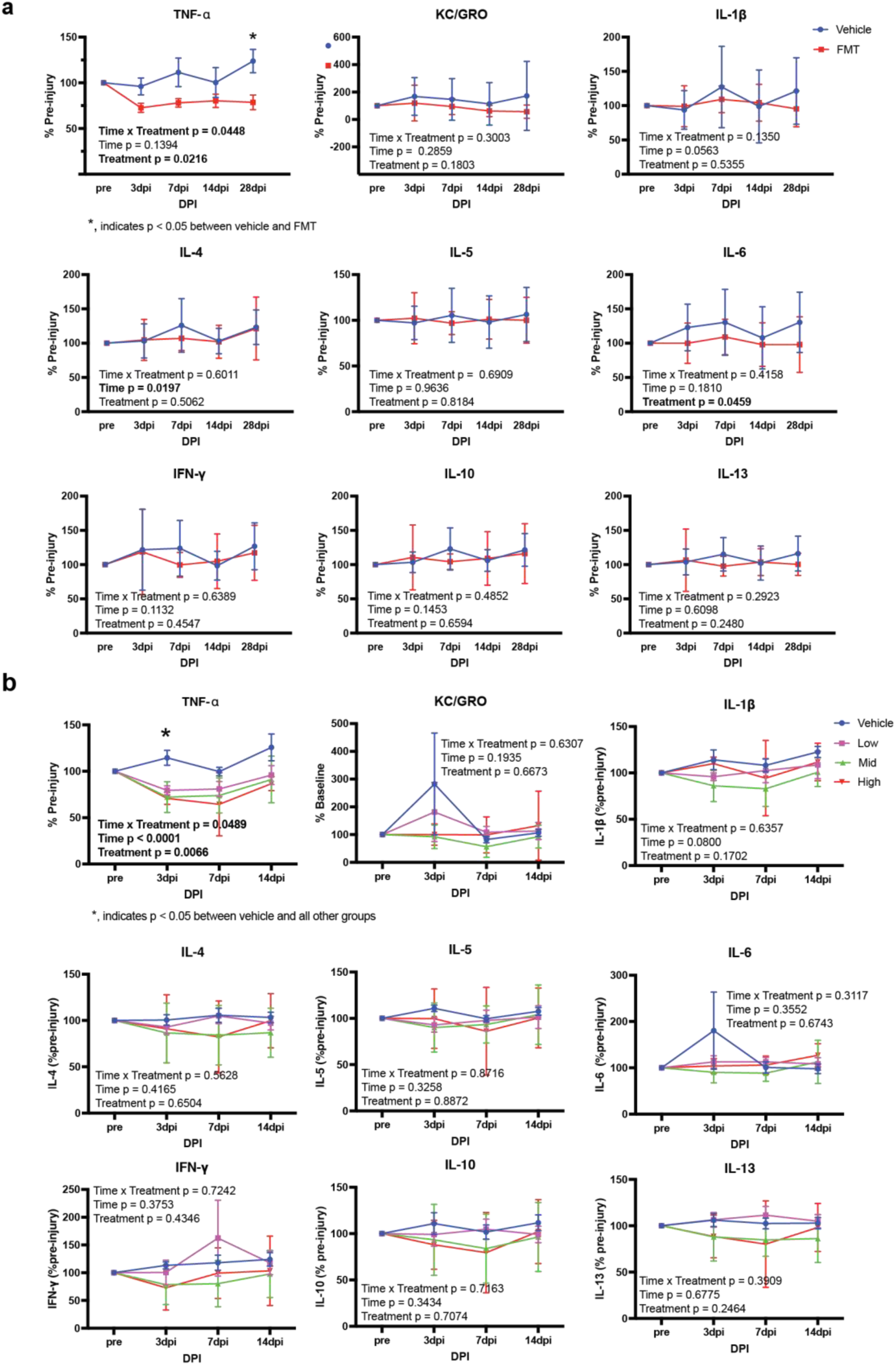
Treating gut dysbiosis reduces the systemic inflammatory response. **a,** Quantification of proinflammatory cytokines in the serum vehicle (blue) and FMT-treated (red) animals pre-injury and at 3, 7, 14, and 28 days post-LC2Hx. Of the nine cytokines tested, TNF-alpha and IL-6 both revealed treatment effects and decreased at all time points post-injury (Two-way ANOVA TNF-alpha p = 0.0216 and IL-6 p = 0.0459). Bonferroni’s posthoc analysis revealed that this decrease was significant in TNF-alpha levels in FMT-treated compared to vehicle-treated animals at 28 dpi (p = 0.0062). **b,** Quantification of proinflammatory cytokines in the serum of vehicle (blue), low-dose (purple), mid-dose (green), or high-dose (red) probiotic animals pre-injury and at 3, 7, and 14 days post-LC2Hx. TNF-alpha revealed a treatment effect and was decreased in all probiotic treated animals compared to vehicle-treated at all timepoints. (Two-way ANOVA TNF-alpha p = 0.0066). Tukey’s posthoc analysis revealed this decrease to be significant at 3dpi in all probiotic-treated groups compared to vehicle (vehicle/low p = 0.0316, vehicle/mid p = 0.0152, vehicle/high p = 0.0139). Bars represent mean ± SD. Bolded text indicates a p-value < 0.05.

We assessed if FMT or probiotic treatment could alter neuronal growth and sprouting characteristics ^49^. To do this, we harvested dorsal root ganglion neurons from the animals in Experiments 2 and 3. Sholl analysis was used to determine the longest neurite length and the total number of crossings or branching of neurites. The summation of neurite length (total outgrowth) and the total number of projections from the soma were also quantified. There were trends of increased total neurite outgrowth, number of crossings, and neurite length in FMT-treated animals, although these changes were not statistically significant (Figures 6 a and b; unpaired, two-tailed t-test; total outgrowth p = 0.3477; t = 0.9445 total number of crossings p = 0.2996; t = 1.056; longest neurite length p = 0.1362; t = 1.532). There was no difference in the number of projections from the soma in FMT versus vehicle-treated animals (Figures 6 b; unpaired, two-tailed t-test; number of projections from soma p = 0.9683; t = 0.03984). Probiotic treatment significantly improved the number of primary projections from the soma, with the high-dose probiotic group producing the greatest number of projections compared to all other groups (Figure 6 a and c; One-way ANOVA p < 0.0001; F = 8.758 with Tukey’s post hoc vehicle/high p < 0.001; low/high p = 0.0002; mid/high p = 0.0067). There was also a trend of increased total neurite outgrowth, and longest neurite length in DRGs treated with high-dose probiotics in comparison to all other groups. Alternately, the mid-dose probiotic group produced the largest total number of crossings (Figure 6 c). These findings indicate that treatment of gut dysbiosis with FMT or probiotics allowed for increased neurite outgrowth potential, longer neurite projections, greater neurite branching ability, and increased sprouting of primary projections from the DRG soma. All of these are potential mechanisms of plasticity and recovery either centrally or in the innervation of the GI tract.

**Figure 6.**
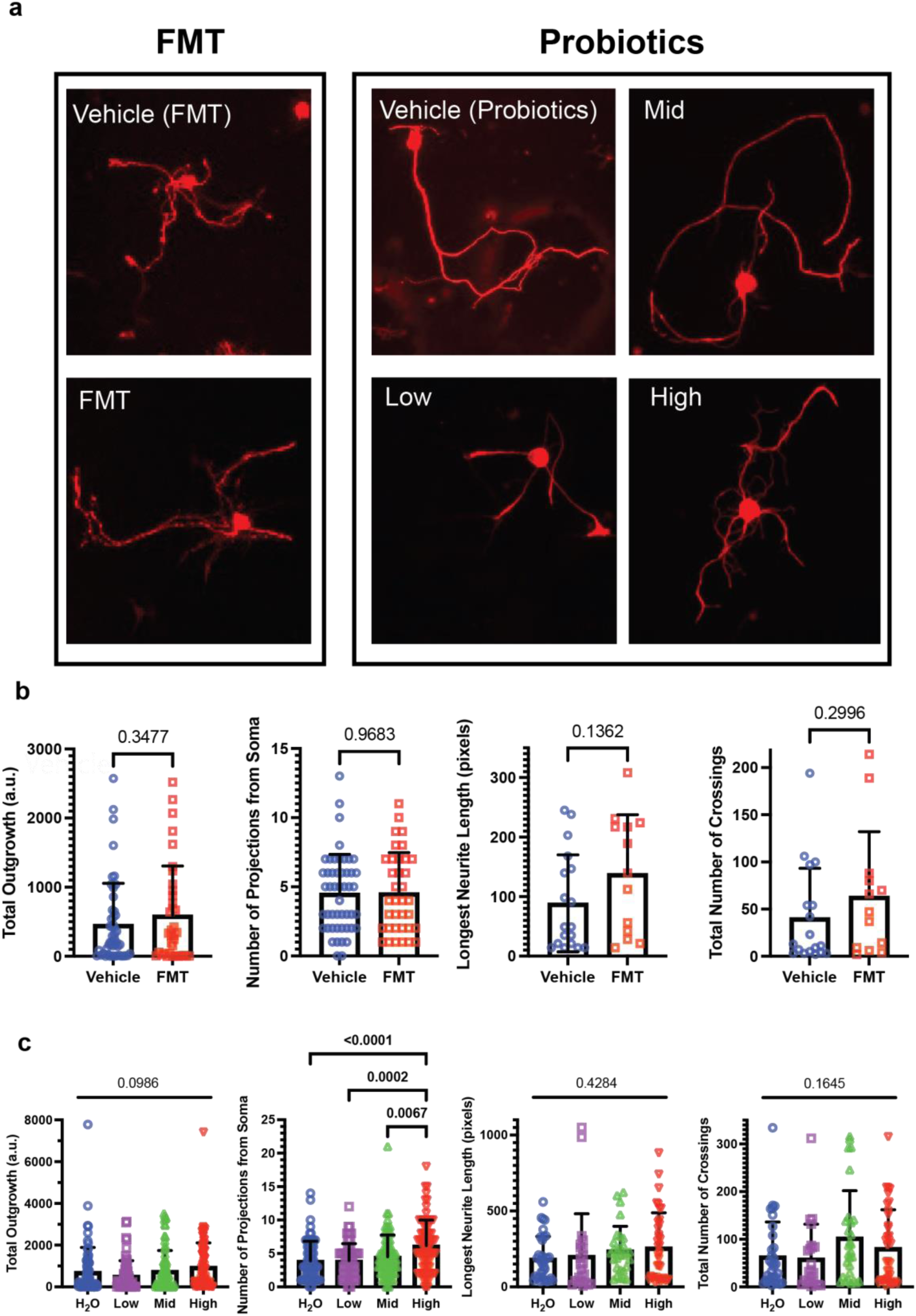
Probiotic treatment following SCI increases the sprouting and regenerative potential of neurons. **a,** Representative images for DRGs treated with vehicle (blue) or FMT (red) (Experiment 2) and vehicle (blue), low-(purple), mid-(green), or high-dose (red) probiotics (Experiment 3). **b,** Mean ± SD of total neurite outgrowth of DRGs *in vitro* measured for *in vivo* FMT versus vehicle-treated animals. There was no significant difference in total outgrowth (p-value = 0.3477). Mean ± SD of the total number of primary projections growing from DRG soma *in vitro* for FMT versus vehicle-treated animals. There was no significant difference in the total number of primary projections growing from the DRG soma (p-value = 0.9683). Mean + SD of the length of the longest measured neurite extending from the DRG. The length of the longest neurite was increased in DRGs from FMT-treated animals (0.1362). Mean + SD of the total number of neurite crossings of the concentric rings in a Scholl analysis of DRGs *in vitro* for animals treated with FMT or vehicle *in vivo*. The total number of crossings was increased in animals treated with FMT (p-value = 0.2996), thus indicating that total neurite branching was increased in animals treated with FMT**. c,** Probiotic treatment did not cause a significant difference in the total neurite outgrowth of DRGs in comparison to vehicle-treated (p-value = 0.0986), although there was a trend towards increased total neurite outgrowth of DRGs from animals treated with high-dose versus low-dose probiotics (p-value = 0.0644, difference = 4.452 +/-1.792). The total number of projections of high-dose probiotics was significantly increased compared to vehicle (2.290 ± 0.5143; p-value < 0.0001****), low-dose probiotics (2.262 ± 0.5351; p-value = 0.0002***), and mid-dose probiotics (1.650 ± 0.5056; p-value = 0.0067**). The length of the longest neurite increased as the concentration of the probiotic treatment increased (p-value = 0.4284). The total number of crossings increased for the DRGs of animals treated with the mid and high concentrations of probiotics (p-value = 0.1645), thus indicating that total neurite branching was increased by the mid and high probiotic concentrations. Bars represent mean ± SD. Bolded text indicates a p-value < 0.05.

Given the enhanced potential to increase primary projections in DRGs in probiotic-treated animals, we next examined the extent of serotonergic sprouting below the level of injury as a potential mechanism for improved respiratory function. Serotonin is critical in activating silent spinal pathways that are important for recovery. We did not observe statistically significant differences in serotonin levels at the C4 ventral horn of animals treated with probiotics versus vehicle at 14 dpi (Supplemental Figure 5 a – b). Because we observed a significant increase in respiratory rate following high-dose probiotic treatment at 7 dpi, examination of serotonergic levels in the brainstem at this more acute time point may be more revealing of a potential mechanism.

## DISCUSSION

Collectively, our experiments present compelling evidence that treating SCI-induced gut dysbiosis improves functional and pathological outcomes. We determined that this dysbiosis is transient, and most severe immediately after LC2Hx. Interestingly, previous studies have determined that interventions for respiratory recovery, including administration of intermittent hypoxia and chondroitinase ABC (ChABC), are more efficacious at chronic time points post-injury ^50–52^. The immediate dysbiosis we observed may be a factor that undermines the effectiveness of early treatment strategies. Understanding the role of the gut-CNS axis and its influence on recovery mechanisms may improve the success of interventions at more acute time points following SCI.

Whole-body plethysmography studies indicate that probiotic treatment improves respiratory rate and overall minute ventilation following cervical SCI. As over half of SCI cases occur at the cervical level, which can impair descending respiratory drive pathways, any improvements in breathing function are of critical importance. Additionally, we found that treatment of gut dysbiosis with probiotics also led to improvements in the pathology of ileal tissue following SCI, demonstrated by an increase in villi length and improvements in the Chiu/Park scoring. Gastrointestinal complications can severely impact an SCI individual’s quality of life, especially those with high-level injuries resulting in tetraplegia. Moreover, 11% of hospitalizations in the SCI population are due to GI complications ^53^. Compromise of gut barrier function can lead to bacterial translocation (leaky gut) and potentially life-threatening septicemia, which is sadly a leading cause of death following SCI. These realities demonstrate the importance of maintaining gut health following SCI for overall well-being and potentially enhancing recovery. This work further emphasizes the importance of personalized and precision medicine through a holistic approach. Studies from our lab have previously demonstrated that genetic background, particularly apolipoprotein E (APOE) genotype, influences respiratory motor plasticity potential ^54^. APOE status has also been robustly associated with distinct microbiome profiles ^55^. Future work evaluating the impact of genotype-associated microbiota composition in injury models may reveal important factors that enhance recovery potential.

Mechanistically, the improvements in respiratory function may be attributed to the reduction in systemic inflammation following the treatment of gut dysbiosis. FMT treatment reduced the systemic IL-6 level following LC2Hx. Additionally, both FMT and probiotic treatment reduced systemic TNF-alpha levels at all time points post-injury compared to vehicle-treated animals. This reduction in systemic proinflammatory cytokine levels may contribute to the observed respiratory recovery, as previous studies have demonstrated that systemic inflammation can impair respiratory plasticity ^41^. Additionally, FMT and probiotic treatment improved the regenerative potential of neurons. The ability for plasticity and regeneration of neurites – either centrally or peripherally - is of utmost importance for SCI recovery. Several mechanisms to improve regenerative capacity have been employed, including enhancing the expression of neurotrophic factors, inhibition of PTEN, administration of intermittent hypoxia, and application of ChABC to digest inhibitory molecules within the glial scar ^51, 52, 56–60^. We now show that treatment of gut dysbiosis can also enhance the regenerative capacity of neurons and can potentially be used in conjunction with these interventions. Overall, these findings provide evidence that treatment of gut dysbiosis following SCI is a critical therapeutic target for recovery at a holistic level.

## Materials and Methods

### Animals and Housing

All experiments were approved by the Institutional Animal Care and Use Committee at the University of Kentucky. Adult female Sprague Dawley rats from Envigo were used in all experiments and allowed to acclimate for at least one week prior to beginning experimental procedures. Animals were co-housed in a 12-hour light/dark cycle with standard chow and water available *ad libitum.* Animals housed together remained with their cage mate throughout the entirety of the experiment and were in the same experimental group to avoid coprophagic cross-colonization. For all experiments, animals were randomized into the specified groups using an online random number generator.

### Groups

The first set of experiments included three cohorts of animals that were separated into three groups: (1) LC2Hx (n =16), (2) surgical sham (n = 12), or (3) drug control (naïve + surgical drugs; n = 8). Animals in the third group received all surgical drugs, including isoflurane anesthesia, and post-surgical care (same as groups one and two) but did not receive any incisions. Group 3 was a necessary control to include because anesthesia and opioids have been shown to impact the gut microbiome ^6, 7^.

The second set of experiments included three cohorts of animals that all received a LC2Hx and were separated into two treatment groups in which they received one of the following types of oral gavage: (1) fecal matter transplant (FMT; n = 12) or (2) vehicle FMT (control; n = 12).

The third set of experiments included two cohorts of animals that all received a LC2Hx and were separated into four treatment groups in which they received one of the following types of oral gavage: (1) vehicle (n = 5), (2) low dose probiotics (n = 6), (3) mid dose probiotics (n = 6), or (4) high dose probiotics (n = 5).

### Fecal Matter Transplant (FMT)

Fecal samples used for the FMT were collected from young (approximately 3 months old), healthy donor animals using an aseptic technique. Immediately after fecal sample collection, samples were pooled and processed to make the FMT slurry solution. Fresh fecal samples were diluted to a 1:10 concentration in sterile PBS (10%), L-cystine HCL (0.05%), glycerol (20%), and sterile water (60%) and filtered with a 100 µm filter to remove fiber content^42^. Vehicle-treated animals received all components of the filtered dilution solution without fecal content. The FMT or vehicle solution was administered via oral gavage for three consecutive days beginning on the day of injury ^42^, and then twice each week until the endpoint at 28 days post-injury ^3^. Each animal received a total of nine oral gavage doses with each dose consisting of 500 µL of the assigned solution.

### Probiotics

The probiotic Visbiome (Lot no. 65026 and 65847) was diluted into the following concentrations: (1) vehicle = sterile water only, (2) low dose = 15 mg (2.7 billion bacteria) in 1 ml sterile water ^43, 44^, (3) mid dose = 130 mg (24 billion bacteria) 1 ml sterile water ^45, 46^, or (4) high = 260 mg (48 billion bacteria) 1 ml sterile water. The assigned concentration of Visbiome was delivered via oral gavage beginning the day of the injury and continued once daily until the endpoint at 14 days post-injury. Each animal received a total of fourteen oral gavage doses, with each dose consisting of 500 µL of the assigned solution.

### Left C2 Hemisection (LC2Hx)

Adult female Sprague Dawley rats were anesthetized with isoflurane gas. Once an appropriate plane of anesthesia was met, animals were prepared for surgery by shaving and sterilizing the surgical site with alternating 70% ethanol and betadine swabs. Ophthalmic ointment was applied to the eyes to prevent drying throughout the procedure. While in the prone position, an incision was made between the ears and extend approximately two inches caudally. The underlying musculature was bluntly dissected and retracted to expose the process of the C2 lamina. The final layer of musculature was then cut and cleared away from the bone so that the intervertebral space between C1 and C2 could be visualized. A C2 laminectomy was performed by cutting the lamina in this space on the left and right sides of the C2 spinous process and extending caudally until the dorsal aspect of the C2 vertebrae was removed and the spinal cord was exposed. Micro-scissors were used to perform a durotomy and a 27-gauge hypodermic needle was bent at a 45-degree angle and used to perform the left C2 hemisection (LC2Hx). Briefly, anatomical landmarks were used to determine midline at the C2 level at which the needle was inserted through the spinal cord until the ventral aspect of bone was reached and then dragged laterally to the animal’s left until completely severing the cord. Three passes were made to ensure complete disruption of pathways on the left side. Finally, all three layers of musculature previously dissected were sutured individually (3-0 absorbable suture) and the skin was closed with surgical staples. All animals received buprenorphine (0.05mg/kg; immediately and 12 hours post-surgery) and carprofen (5mg/kg; immediately, 12-, and 24 hours post-surgery) subcutaneously. Surgical animals were monitored twice daily for the four days post-injury with subcutaneous saline given as needed. Animals recovered for one week on a heated rack separated from naïve animals and remained on a standard 12-hour light/dark cycle with food and water accessible *ad libitum* prior to returning to standard housing. Welfare and weights were checked weekly throughout the entire experiment.

### Whole Body Plethysmography (WBP)

Whole Body Plethysmography (WBP) (DSI, Buxco FinePointe) was used to assess ventilatory patterns in awake, unanesthetized rats before and after LC2Hx. At the designated timepoints pre-and post-injury, animals were placed into individual DSI whole body plethysmography chambers and breathing measurements were recorded every two seconds for the duration of approximately one hour. At the beginning of each experiment, an acclimation period of 20-minutes under normoxic conditions (21% O_2_) was used to minimize atypical respiratory patterns while initially exploring the chamber (i.e., sniffing). Immediately after, a 30-minute baseline recording was taken under normoxic conditions (21% O_2_), followed by a 10-minute hypoxic challenge (11% O_2_), and finally a 10-minute post-hypoxia recording under normoxic conditions (21% O_2_). From the recordings, we examined respiratory rate (breaths/min), tidal volume (mL/breath/g), and minute ventilation (mL/min/g). Food and water were restricted while breathing measurements were taken. At the conclusion of the experiment, animals were returned to their home cages where food and water were accessible *ad libitum*.

### Diaphragmatic Electromyogram (diaEMG) Recording

Animals were anesthetized with isoflurane gas using the SomnoSuite low-flow anesthesia system (Kent Scientific). Once an appropriate level of anesthesia was met, animals were transferred into a faraday cage and switched to a nose cone for continued delivery of approximately 2.4% isoflurane throughout the procedure. Animals were placed in the supine position, and a laparotomy was performed. Upon visualization of the phrenic nerve on the diaphragm, bipolar needle electrodes were inserted bilaterally into the dorsolateral quadrant of the costal hemidiaphragm. Electrodes were placed in close proximity to the phrenic nerve while being cautious to avoid piercing the nerve. Bipolar electrodes were connected to an amplifier and data acquisition system (BMA-400 Four-channel Bioamplifier, CWE, Inc., Ardmore, PA, USA; CED 1401 with Spike2 Data Analysis Computer Interface, CED, Cambridge, UK) to record and analyze the electromyogram (EMG) signal. Following election insertion, the faraday cage was closed, and a 10-minute baseline recording was measured. Immediately following the baseline recording, three 15-second nasal occasions were performed to assess each animal’s maximal respiratory output. Sufficient time between nasal occlusions was allowed for burst amplitude to return to approximately pre-occlusion levels. At the conclusion of the EMG recording, animals were given ketamine/xylazine intraperitoneally while being weaned off isoflurane and transitioned to lung health experiments.

### 16S rRNA Sequencing

Fecal samples were collected from study animals using an aseptic technique pre-and at various timepoints post-LC2Hx in order to assess the gut microbiome. Samples were immediately stored at −80°C until all samples were collected for further processing. The bacterial 16S rRNA gene was then isolated from the fecal samples using DNeasy PowerSoil Pro Kit (Ref #47014 and 47016; Lot #169034368 and 17203302). DNA from each sample was diluted to a concentration ranging between 1-50ng/µL and mailed to Argonne National Laboratory for sequencing.

*Read processing*: 16SV4 was amplified using 515F-806R primers. Fastq files were demultiplexed using qiime2 demux emp-paired function. Raw data were processed with qiime2. Briefly, reads were denoised with dada trimming nucleotides based on quality plots (trim reads when the average quality <30-see code for details). Unique reads were clustered to 99% identity using the silva-138-99-seqs-515-806 database (ASV99% or OTUs). Chimeras were removed using vsearch uchime-denovo function. Samples had an average of 29,245 reads (Table 1). ASVs with <50 reads total abundance were removed since they were present in <2 samples and will not significantly contribute to the analysis. ASVs were classified using the feature-classifier classify-sklearn function with the silva-138-99-515-806-nb-classifier. When taxonomic classification was not available at lower levels, the immediately following taxonomic level classification was used. Filtered ASV 99% table (referred to as OTUs in the figures) was subsampled to analyze 15,000 reads per sample. Two samples with an unusually low number of reads were removed.

**Table 0.1.** Summary of 16S sequencing reads processing.

|  | **Number**<br>**(mean)** | **% of reads** |
| --- | --- | --- |
| **Raw input (91 samples)** | 29245.0889 | 88.55 |
| **Denoised dada** | 25398.6111 | 86.77 |
| **ASV 100% (total)** | 2305 | NA |
| **ASV 99% (total)** | 1707 | NA |
| **ASV 99% (per individual)** |  |  |
| **ASV 99% non-chim** | 1,393 | NA |
| **ASV 99% non-chim (per individual)** |  |  |
| **ASV 99% non-chm filtered (>50 reads, >n=2; per individual)** | 896 | NA |
| **ASV 99% non-chm filtered (>50 reads, >n=2; per individual)** | 896 | NA |
| **ASV 99% non-chm filtered rarefied (15,000 reads; 89/91 samples)** | 896 | NA |
| **Taxonomic levels (Family) from ASV99% non-chm filtered rarefied** | 48 | NA |

#### Read processing

OTU abundance matrix was analyzed using vegan, and labdsv packages in R. Principal coordinate analysis was performed on rarefied reads and log10 normalized reads at the OTU and family taxonomic level. Simper analyses were performed to identify OTUs that explain most of the variation in the ordination space.

#### Statistical analysis

Statistical analysis of group separation in the ordination space was performed calling Permanova with the Adonis function in R. Mixed-effect time series statistical analysis of clusters and OTUs abundance was done using prism.

#### Sectioning and Staining CNS

Following euthanasia at the designated times post-injury, the skull and spinal column were removed and drop-fixed in 4% paraformaldehyde (PFA) for 48 hours at 4°C. The brainstem and cervical spinal cord were then dissected from the skull and spinal column and placed in fresh 4% PFA for an additional 24 hours at 4° and then cryoprotected in 30% sucrose at 4°C until sectioning. The injury was used as the epicenter to create three 4mm sections (brainstem, C1/2, and C3/4 levels). Tissue sections were then embedded in Optimal Cutting Temperature embedding media (OCT, Sakura Tissue-TEK), and cryosections were taken at 30µm (Leica cryostat) and placed directly onto gelatin-coated slides (Thermo Fischer Scientific) and stored at-20°C.

#### Cresyl Violet

Slides were placed in CitriSolv solution prior to serially rehydrating the tissue in decreasing concentrations of ethanol. Slides were then rinsed in distilled water, stained with 0.5% Cresyl Violet solution (Sigma Cat #C5042), rinsed in distilled water, dehydrated in the reverse order of ethanol concentrations used, and placed in CistroSolv. Coverslips were then applied using permount (Electron Microscopy Sciences Cat #17986-01). Slides were imaged on a Keyence BZ-X810 microscope (Keyence Corporation of America, Itasca, IL, USA) using bright field illumination at 2x.

#### Immunohistochemistry (IHC)

Frozen sections were thawed to room temperature (approximately 20°C) and placed in Heat-Induced Epitope Retrieval at 85°C for 15 minutes. Next, slides were permeabilized with 0.4% TritonX-100 in 1xPBS for 15 minutes prior to blocking in a solution of 3% normal goat serum, 0.2% TritonX-100 in 1xPBS for 60 minutes at room temperature. Slides were then incubated in 5-HT primary antibody diluted to 1:2000 (rabbit, ImmunoStar Cat #20080; Lot #1650001) and NeuN primary antibody diluted to 1:5000 (mouse, ImmunoStar Cat #MCA-1B7; Lot #080522) at 4°C overnight. Next, slides were washed with 1xPBS before incubating in goat anti-rabbit AlexaFluor 594 (Life Technologies Ref #A11037; Lot #1777945) and goat anti-mouse IgG AlexaFluor 488 (Life Technologies Ref #A11029; Lot #1745855) for 120 minutes at room temperature. Finally, slides received a final wash in 1xPBS and were mounted with Fluoromount-G (SouthernBiotech Cat #0100-01). Slides were imaged on a Keyence BZ-X810 and analyzed using HALO v2.3.2089.70 (Indica Labs, Inc.).

#### Gut Tissue (Swiss Roll)

Following euthanasia at the designated times post-injury, the laparotomy incision made for the diaEMG recording was expanded distally to fully visualize the abdominal cavity. The stomach and cecum were then identified as markers for the beginning and end of the small intestine. The small intestine was then removed and divided into thirds, with the final third known as the ileum. The ileum was then flushed with 1xPBS and rolled into a swiss roll configuration for sectioning as previously described ^61, 62^. In brief, the ileum was cut longitudinally along the mesentery attachment to expose the lumen. A 1ml syringe plunger was used to roll the ileum from the distal to the proximal end in the formation of the swiss roll. The tissue was then placed in 4% PFA at 4°C for 24 hours and then moved to 70% EtOH until paraffin embedding and sectioning at a thickness of 5µm. Slides were stained with hematoxylin and eosin (H&E) and Alcian blue for histological assessment and summation of goblet cells respectively. To assess intestinal injury, the Chiu/Park scoring system was employed and all evaluators were blinded to injury or treatment. The Chiu/Park score ranges on a scale of 0 – 5 (0 = normal mucosa; 1 = subepithelial space at the villous tip; 2 = extension of subepithelial space with moderate lifting; 3 – massive lifting down the sides of villi, some denuded tips; 4 = denuded villi, dilated capillaries; 5 = disintegration of lamina propria) ^20^.

Additionally, slides underwent MALDI imaging. A Waters Synapt G2Si mass spectrometer (Waters Corporation, Milford, MA) equipped with an Nd:YAG UV laser with a spot size of 50um was used to detect released glycogen. A total of 300 laser spots per pixel at X and Y coordinates were collected at a raster size of 100um. Data acquisition, spectrums were uploaded to High-Definition Imaging (HDI) Software (Waters Corporation) for mass range analysis from 750 to 4000m/z. HDI generated glycogen images were obtained.

#### Systemic Inflammation

Blood was collected via the lateral tail vein at the designated timepoints pre-and post-injury. Serum levels of IFN-γ, IL-1β, IL-4, IL-5, IL-6, IL-10, IL-13, KC/GRO, TNF-α were assayed using the Meso Scale Discovery (MSD) V-PLEX Proinflammatory Panel 2 Rat Kit (Lot #K002026; Catalog #K15059D-1; Rockville, Maryland 20850-3173).

#### Lung Function Measurements

Rats were anesthetized by intraperitoneal injection of xylazine and ketamine. After a tracheotomy, the rats were connected to the flexiVent system (scireq, an emka technologies company, Montreal, QC Canada H2S 3G8). The computer-controlled small animal instrument ventilated the rats with a tidal volume of 10 ml/kg at a frequency of 90 breaths/minute. Two snapshots, two total lung capacity maneuvers, three more snapshots, and three pressure-volume loop perturbations were performed for each individual subject on the flexiVent. From the second set of snapshots after full inflation, the three acceptable recorded measurements were averaged for dynamic compliance and dynamic resistance. An estimate of inspiratory capacity was averaged from the three total lung capacity maneuvers.

Bronchioalveolar lavage (BAL) fluid was collected using two 5mL syringes loaded with 4mL BAL fluid buffer (sterile PBS + 0.1% EDTA). Following tracheotomy and flexiVent, the loaded syringe was connected to the cannula and the fluid was slowly injected, then slowly retrieved to collect BAL fluid in the same syringe. This process was repeated with the remaining syringe. Aliquots of BAL fluid were combined in a 15mL conical tube and spun at 200 x g at 4C for 10 minutes. Three 500uL aliquots of BAL fluid were recovered for future processing and the remaining supernatant was discarded. The pellet was resuspended in 500uL PBS and placed on cytospin slides (30,000 cells/slide) for differential cell counting. The following calculation was used to determine cells/mL BAL: (Live cells (cells/mL) * volume of reconstituted pellet (0.5mL)) / (volume of BAL recovered) = cells/mL BAL.

#### Open Field Chamber

At the designated timepoints pre-and post-injury, rats were placed in SuperFlex Open Field System chambers (16×16 inches; Omnitech Electronics Inc., Colubus, OH, USA) for five minutes and allowed to explore freely while lateral and horizonal movements were recorded. Total distance, vertical activity count, vertical time, and center time were measured using Fusion Software v6.5 and analyzed using GraphPad Prism v9.5.0.

#### Dorsal Root Ganglia (DRG) Outgrowth

DRGs were harvested at 28 DPI for FMT-treated animals, and 14 DPI for probiotic-treated animals. After excising the spinal column from the animal, it was bisected along the dorsal and ventral aspects to expose the DRGs. Those were then removed and cleaned of any peripheral projections. Cleaned DRGs were then incubated with enzymes (2000U/mL collagenase: 50U/mL dispase) overnight to digest and dissociate individual cell bodies.

Dissociated DRGs were cultured on a substratum of 0.1mg/mL Poly-D-Lysine (Life Technologies) and 5ug/mL Laminin (Life Technologies) with Serum Free Media (Neurobasal A media, B27 Supplement, Penicillin-Streptomycin, GlutaMAX; Life Technologies). Media was exchanged at 24 hours to contain anti-mitotic agents Fluorodeoxyuridine (4mM) and Cytosine Arabinoside (100LM) to limit satellite cell contamination. Cells were allowed to grow for 3-5 days based on visible confluency, were then fixed with 4% paraformaldehyde and stained with Beta-III Tubulin to visualize neurons for analysis.

A Keyence BZ-X810 Fluorescence Microscope was used to image take 24 random images per animal. Image locations were chosen randomly. All cells within each image for which the entirety of the cell outgrowth was visible was quantified via ImageJ software utilizing the NeuronJ plug-in. This quantification includes the total neurite outgrowth, total number of primary projections originating from the soma, the length of the longest neurite (Sholl analysis), and total neurite branching (Sholl analysis).

## Statistical Analysis

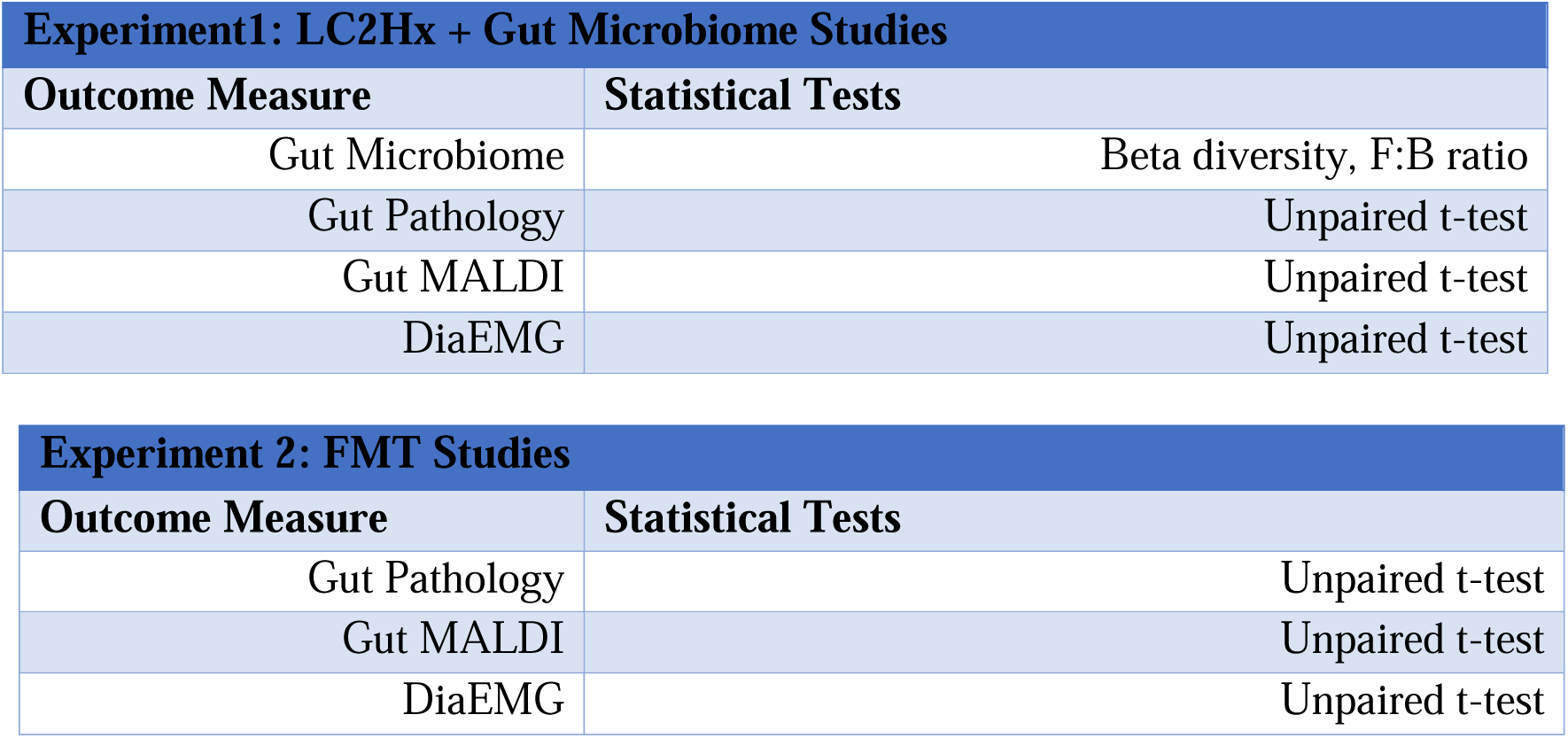
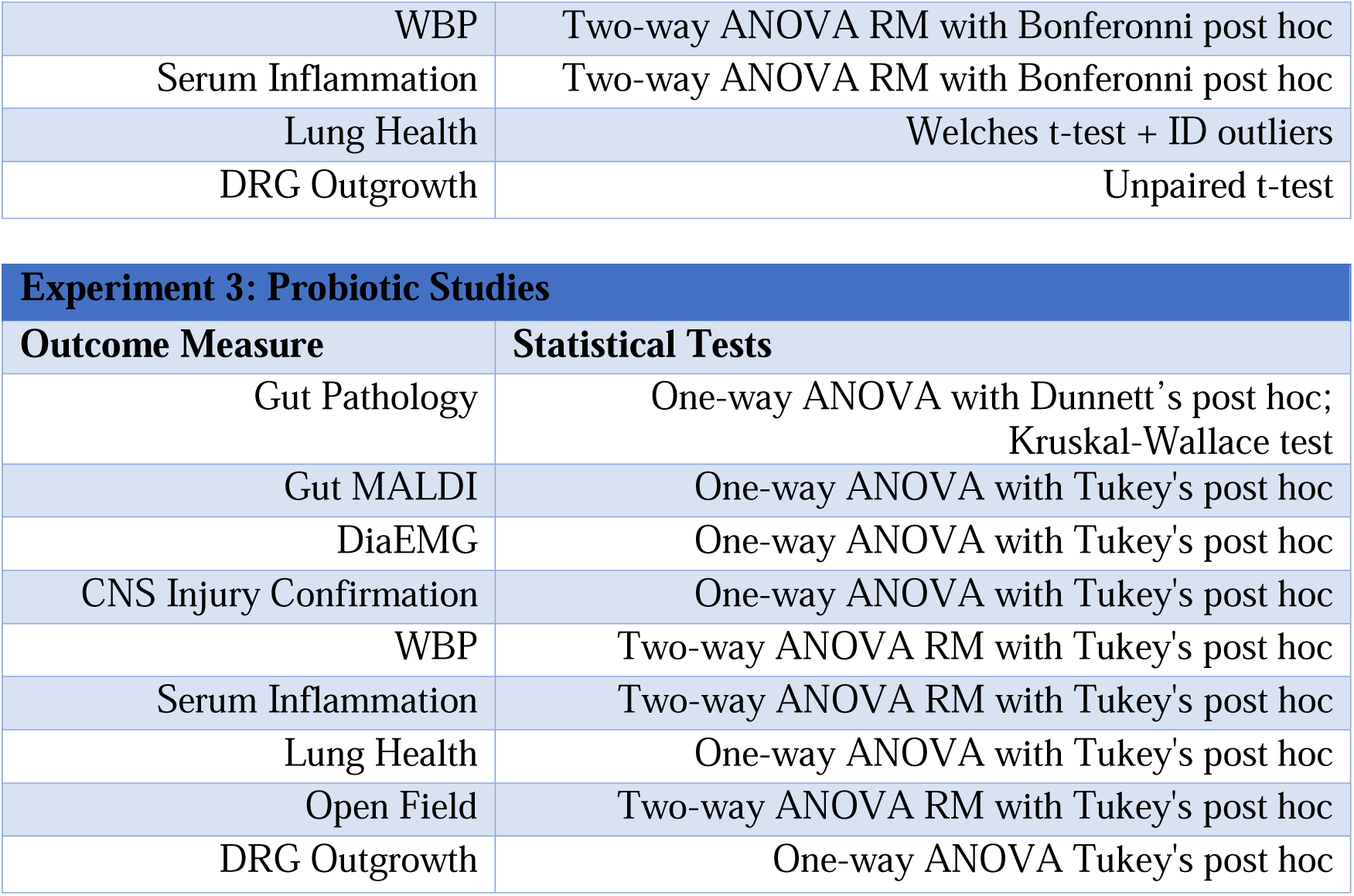

## ACKNOWLEDGEMENTS

This study was supported by funding through:

R01 NS101105 (WJA) and NIH R21 NS121966 (WJA)

NIH Training T32 NS077889 (JSN), Neurobiology of CNS Injury and Repair

Kentucky Spinal Cord and Head Injury Research Trust (KSCHIRT)

R35NS111582 (PGP and KAK) and Ray W. Poppleton Chair (PGP)

NIH grants HL151419 and HL131526, and the Kentucky Research Alliance for Lung Disease (CMW)

R01AG066653, R01CA266004, R01AG078702, CureAlz fund (RCS)

Research reported in this publication was supported by an Institutional Development Award (IDeA) from the National Institute of General Medical Sciences of the National Institutes of Health under grant number P30 GM127211.

## AUTHOR CONTRIBUTIONS

J.N.W. contributed to the conception and design of experiments, acquisition, analysis, and interpretation of data, the initial draft of the work, and revisions.

K.A.K experimental design, analysis of 16S rRNA data, and draft revision

M.D.S. experimental design, acquisition, and analysis of data, and draft revision

C.E.T. experimental design, acquisition, and analysis of whole-body plethysmography (WBP) data, draft revision

S.L.S. acquisition, analysis, and interpretation of dorsal root ganglia (DRG) data, analysis of ileal tissue data

B.C.R. acquisition and analysis of ileal tissue data

C.M.C. experimental design, acquisition, and analysis of data, draft revision

B.E.D. acquisition and analysis of lung function data

T.R.H. acquisition and analysis of Matrix-Assisted Laser Desorption and Ionization (MALDI) data

H.A.C. acquisition of Matrix-Assisted Laser Desorption and Ionization (MALDI) data

A.D.B. acquisition, analysis, and interpretation of data and review of the manuscript

C.M.W conception and design of experiments, data interpretation, and review of the manuscript

R.C.S. conception and design of experiments, data interpretation, and review of the manuscript

P.G.P. conception and design of experiments, data interpretation, and review of the manuscript

W.J.A. conception and design of experiments, acquisition, analysis, and interpretation of data, initial draft of the work, and revisions

---

**Supplemental Figure 1.**
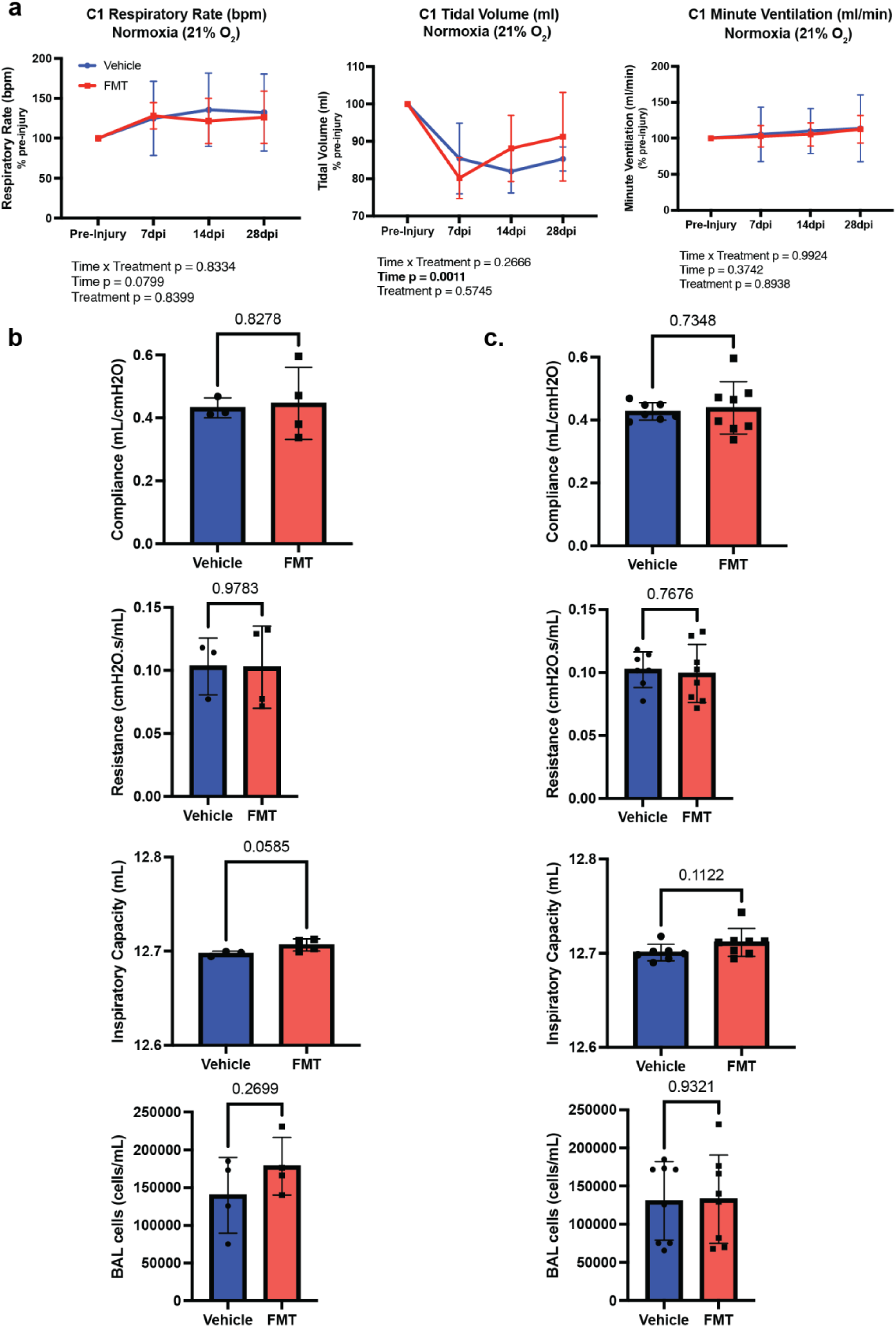
FMT treatment does not improve ventilatory function following LC2Hx. **a,** Quantification of WBP measurements (respiratory rate, tidal volume, and minute ventilation) in cohort 1 (n = 4 animals/group) following LC2Hx and treatment with vehicle (blue) or FMT (red) revealed marginal differences between groups, although there appeared to be some improvement in the tidal volume of FMT-treated animals at 14 and 28 dpi. (Two-way ANOVA treatment p = 0.5745). **b,** Lung function measurements (compliance, resistance, inspiratory capacity, and BAL cell counts) in cohort 1 (n = 4 animals/group) revealed no significant differences between groups (Welch’s t-test compliance p = 0.8278, resistance p = 0.9783, inspiratory capacity p = 0.0585, BAL cell counts p = 0.2699). **c,** Lung function measurements (compliance, resistance, inspiratory capacity, and BAL cell counts) in cohorts 1 and 2 (n = 8 animals/group) revealed no significant differences between groups (Welch’s t-test compliance p = 0.7348, resistance p = 0.7676, inspiratory capacity p = 0.1122, BAL cell counts p = 0.9321). Bars represent mean ± SD. Bolded text indicates a p-value < 0.05.

**Supplemental Figure 2.**
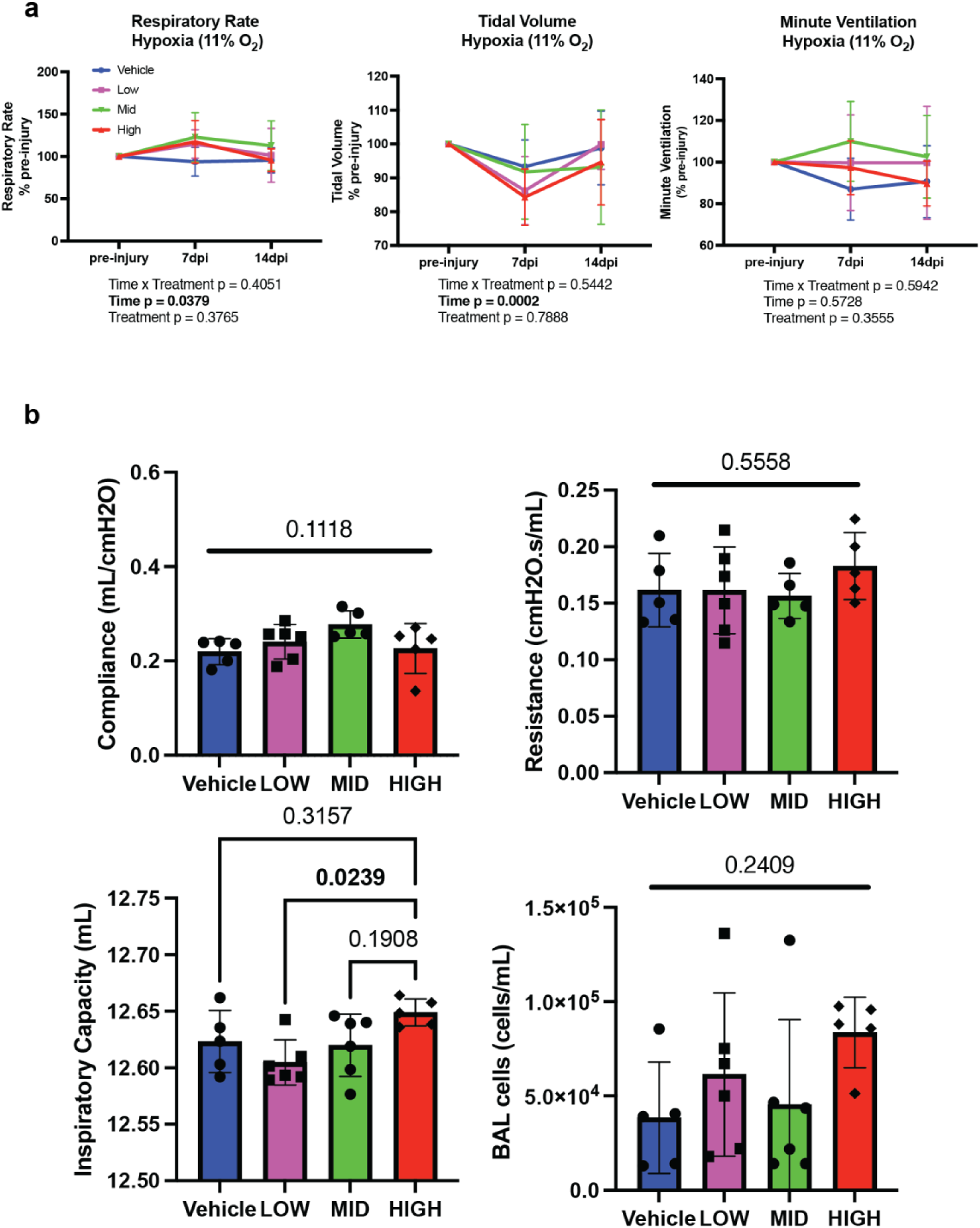
Hypoxic whole-body plethysmography and lung mechanic measurements following SCI and probiotic treatment. **a,** Quantification of WBP measurement (respiratory rate, tidal volume, and minute ventilation) under hypoxic conditions (11% oxygen) in vehicle (blue), low-dose (purple), mid-dose (green), or high-dose (red) probiotic animals pre-injury, 7, and 14 days post-LC2Hx revealed marginal differences between groups. There appeared to be some improvement in the minute ventilation of probiotic-treated animals at 7 dpi, as all probiotic-treated groups produced an increase in minute ventilation compared to vehicle, although this increase was not statistically significant (Two-way ANOVA treatment p = 0.3555). **b,** Lung function measurements revealed no significant differences in compliance, resistance, or BAL cell counts (One-way ANOVA compliance p = 0.1118, resistance p = 0.5558, BAL cell counts p = 0.2409). Inspiratory capacity differed between groups, with the high-dose group significantly higher than the low-dose group (One-way ANOVA inspiratory capacity p = 0.0585, Tukey’s posthoc low/high p = 0.0239). Bars represent mean ± SD. Bolded text indicates a p-value < 0.05.

**Supplemental Figure 3.**
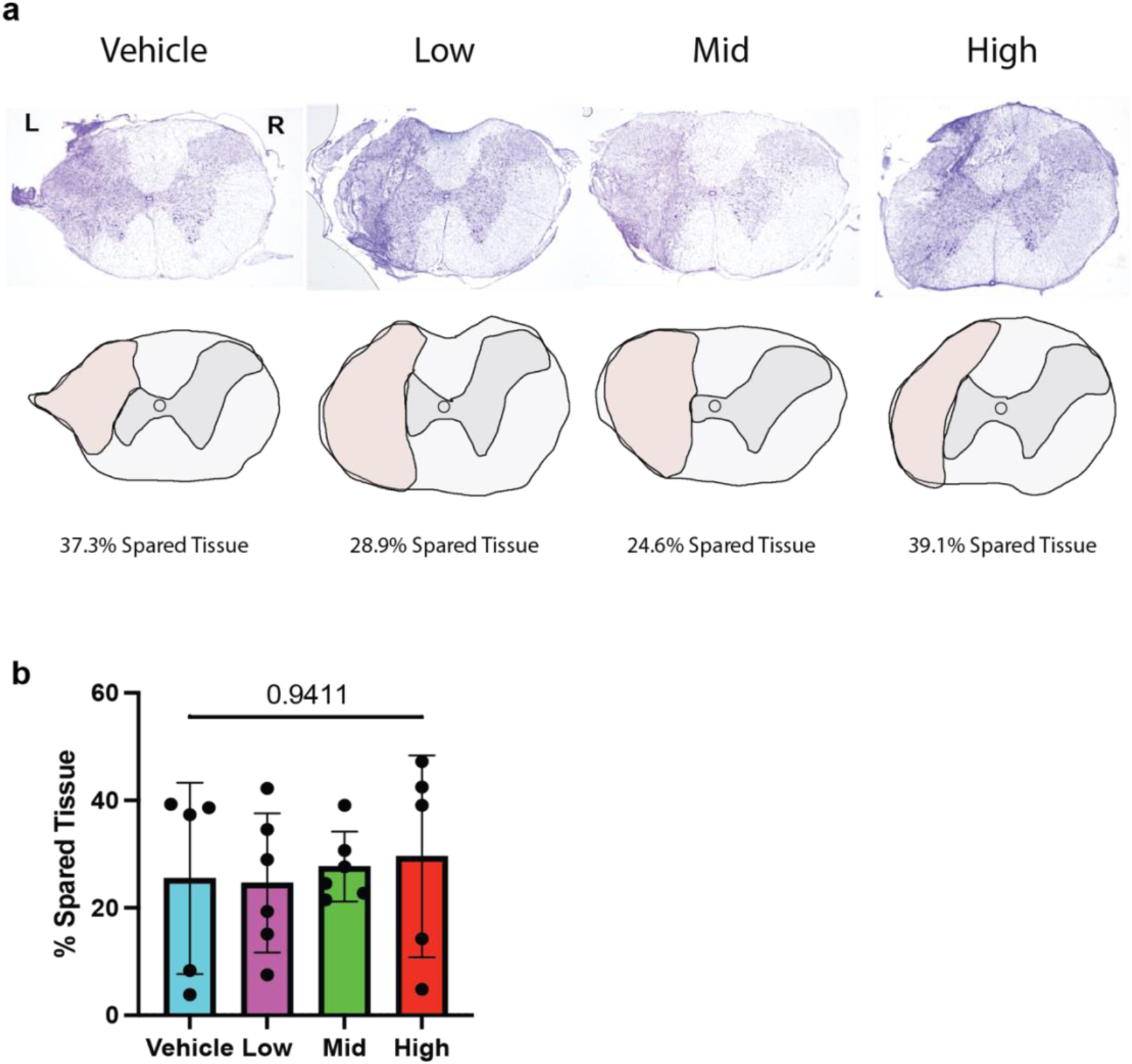
Confirmation of LC2Hx in Experiment 3: Probiotic treatment following LC2Hx. **a,** Representative images of cresyl violet staining at the level of injury (spinal level C2) in vehicle, low-dose, mid-dose, and high-dose probiotic-treated animals 14 days post-LC2Hx. **b,** Quantification of spared tissue indicated no significant differences between groups (vehicle = blue, low = purple, mid = green, high = red) at the level of injury, demonstrating that all animals were equally injured (One-way ANOVA p = 0.9411). Therefore, improvements observed in outcome measures are not attributed to a less severe injury. Bars represent mean ± SD.

**Supplemental Figure 4.**
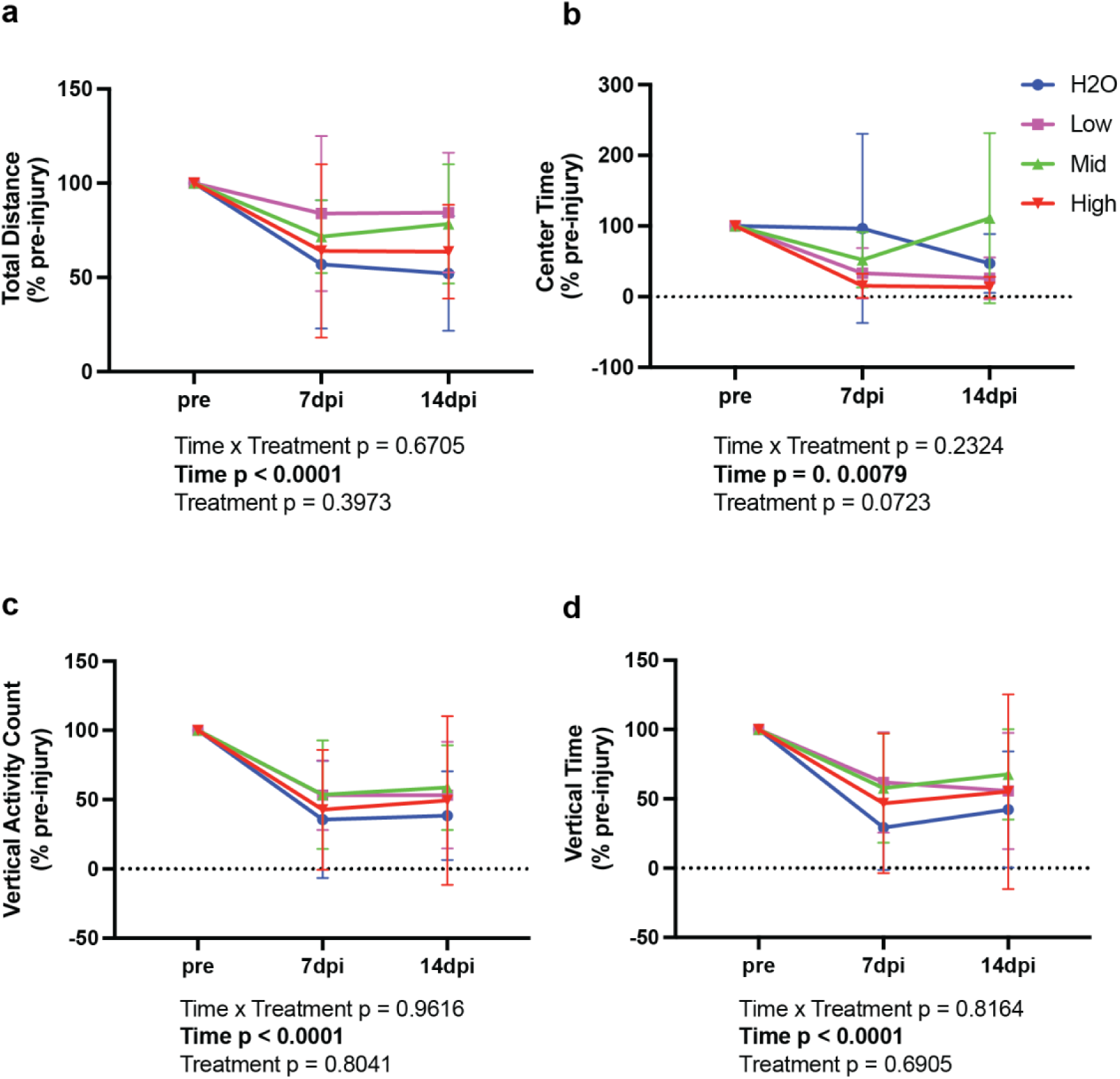
Probiotics treatment does not affect locomotor activity following LC2Hx. Evaluation of locomotion was assessed in activity chambers pre-injury and at 7 and 14 dpi. **a,** All probiotic-treated groups demonstrated an increase in total distance traveled at both time points post-injury, although this trend was not statistically significant (Two-way ANOVA treatment p = 0.3973). **b,** Center time was higher in vehicle-treated animals at 7 dpi compared to all probiotic groups, although not significantly (Two-way ANOVA treatment p = 0.0723). Bonferroni post hoc analysis revealed no significant differences between groups at any time point. **c – d,** There were no significant differences between groups in **c,** vertical activity count, or **d,** vertical time (Two-way ANOVA vertical activity count p = 9616, vertical time p = 8164). As expected, there was a time effect in all measurements, as all animals were injured. Bars represent mean ± SD. Bolded text indicates a p-value < 0.05

**Supplemental Figure 5.**
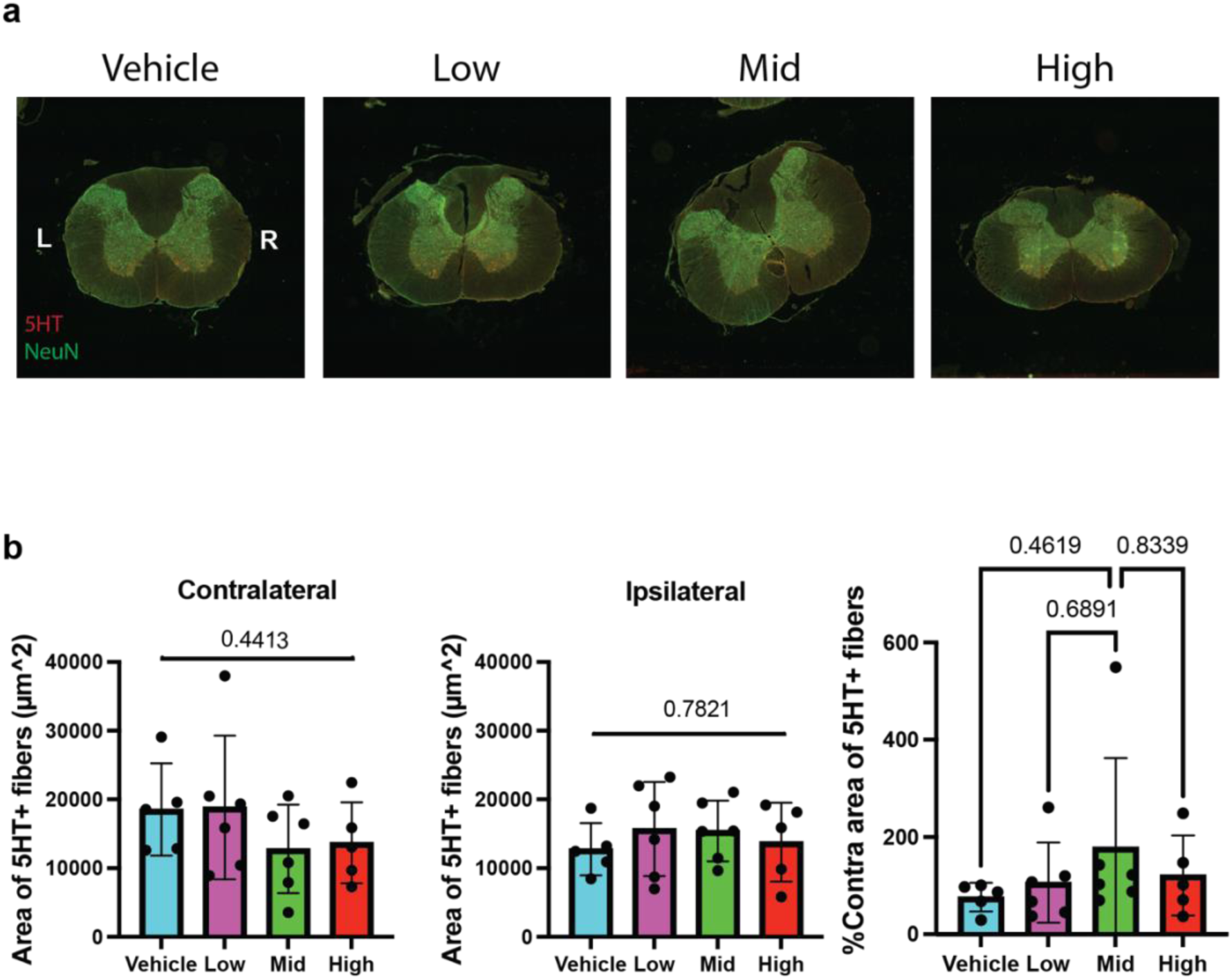
Probiotic treatment does not alter serotonin levels at the level of the phrenic motor nucleus. **a,** Representative images of 5-HT (serotonin) staining at the level of the phrenic motor nucleus (spinal level C4) in vehicle, low-dose, mid-dose, and, high-dose probiotic-treated animals 14 days post-LC2Hx (5-HT = red, NeuN = green). **b,** Quantification of 5-HT staining indicated no significant differences between groups (vehicle = blue, low = purple, mid = green, high = red) at the level of C4 on the contralateral or ipsilateral side of injury (One-way ANOVA contralateral p = 0.4413, ipsilateral p = 0.7821). Evaluation of the ipsilateral side as a percentage of the contralateral side indicated that all probiotic groups had more serotonin than vehicle-treated, although this trend was not statistically significant (One-way ANOVA p = 0.6941). Bars represent mean ± SD.

